# Symbiotic diazotrophic UCYN-A strains co-occurred with El Niño, relaxed upwelling, and varied eukaryotes over 10 years off Southern California Bight

**DOI:** 10.1101/2022.11.07.514914

**Authors:** Colette Fletcher-Hoppe, Yi-Chun Yeh, Yubin Raut, J.L. Weissman, Jed A. Fuhrman

**Affiliations:** Marine & Environmental Biology, Dornsife College of Letters, Arts and Sciences, University of Southern California (USC); Department of Biological Sciences, Dornsife College of Letters, Arts and Sciences, University of Southern California (USC)

## Abstract

Biological nitrogen fixation, the conversion of N2 gas into a more bioavailable form, is vital to sustaining marine primary production. Studies have shifted beyond traditionally studied tropical diazotrophs. *Candidatus* Atelocyanobacterium thalassa (or UCYN-A) has emerged as a research focal point due to its streamlined metabolism, intimate partnership with a haptophyte host, and broad distribution. Here, we explore the abiotic factors that govern UCYN-A’s presence at the San Pedro Ocean Time-series (SPOT), its partner fidelity, and statistical interactions with non-symbiotic eukaryotes. 16S and 18S rRNA sequences were amplified by “universal primers” from monthly samples and resolved into Amplicon Sequence Variants, allowing us to observe multiple UCYN-A symbioses. UCYN-A1 relative abundances increased following the 2015-2016 El Niño event. When this “open ocean ecotype” was present, coastal upwelling ceased, and Ekman transport brought tropical waters into the region. Network analyses reveal all strains of UCYN-A co-occur with dinoflagellates including *Lepidodinium*, a potential predator, and parasitic *Syndiniales*. UCYN-A2 appeared to pair with multiple hosts and was not tightly coupled to its predominate host, while UCYN-A1 maintained a strong host-symbiont relationship. These biological relationships are particularly important to study in the context of climate change, which will alter UCYN-A distribution patterns both locally and globally.

## Introduction

Biological nitrogen fixation sustains primary production in much of the ocean. In this process, rare prokaryotes known as diazotrophs convert inert dinitrogen gas (N_2_) to ammonia (NH_3_). Only a handful of well-characterized photosynthetic bacteria were once known as diazotrophs: *Trichodesmium, Crocospharea watsonii*, and symbionts in diatom-diazotroph associations (DDAs) were considered the dominant diazotrophs (e.g. [1, 2]). However, traditional paradigms of biological nitrogen fixation are continuously being challenged (e.g. [2]). Many studies have applied amplicon sequencing to target the gene *nifH*, which encodes a subunit of the nitrogenase enzyme that conducts nitrogen fixation, and revealed that diazotrophs are a more diverse group than previously recognized (e.g. [3–6]). The first study that applied this technique to marine organisms observed a cluster of *nifH* sequences belonging to a clade termed “UCYN-A”, for “Unicellular Cyanobacterium A” [7]. This clade of organisms has been tentatively named *Candidatus* Atelocyanobacterium thalassa [8], and is now recognized as a major contributor to biological nitrogen fixation (e.g. [9, 10]).

UCYN-A is an aberrant cyanobacterium, lacking photosystem II (PSII) and key components of cellular pathways, such as the Krebs cycle [11]. Its metabolism is streamlined because it lives in symbiosis with a photosynthetic haptophyte host, exchanging fixed nitrogen for carbon compounds [8, 12]. Four clades of UCYN-A are currently recognized based on their *nifH* sequences, although more may exist [13]. UCYN-A1, the most extensively studied type of UCYN-A, associates with a coccolithophore member of the genus *Braarudosphaera* [8], and is found primarily in open-ocean regions [14, 15]. UCYN-A1 is <1μm in diameter while host cells have a diameter of 1-3μm and can house 1-2 symbionts each [13, 16]. UCYN-A2, a coastal ecotype [14, 15], associates with the coccolithophore *Braarudosphaera bigelowii* [12, 17], and is larger than 1μm, while host cells are 4-10μm in diameter (e.g. [18]) and can house 4-10 symbionts per cell [14, 16]. UCYN-A1 and UCYN-A2 are thought to have diverged from one another ~91 million years ago [19].

UCYN-A has a broad, global distribution [13], including nitrogen-rich systems, such as coastal and equatorial upwelling systems (e.g. [2, 10, 20]). UCYN-A has been found previously in the Southern California Current System and Monterey Bay [21, 22]. Notably, UCYN-A has been found at our study site, the San Pedro Ocean Time-series (SPOT) as deep as 890m [23], and at a nearby daily time series off the coast of Catalina Island [24]. Furthermore, UCYN-A was found to comprise up to 95% of the diazotroph population sampled from San Diego to Sebastian Vizcaino Bay (Baja, CA) and within the period of our 10-year timeseries [21].

Nitrogen fixation by many species has been observed in coastal ecosystems in surprisingly high rates, often despite high concentrations of organic nitrogen. For example, nitrogen fixation off the New Jersey Shore may support up to 100% of primary production in this ecosystem, with some of the highest reported UCYN-A abundances [25]. Although coastal nitrogen fixation is speculated to be quite high, it may only be detected at low rates due to eukaryotic grazing that rapidly consumes diazotrophs as they fix nitrogen [10]. However, few studies have attempted to link diazotrophs to potential predators, and these reports have focused on diazotrophs confined to the tropics [26, 27].

In addition, many questions remain about the UCYN-A symbiosis, including the specificity of host-symbiont partnerships (e.g., UCYN-A has been reported without a host [19]). The question of UCYN-A host specificity is especially important because its established hosts are under threat from climate change and ocean acidification, which may degrade the calcareous shells of these coccolithophores [28]. This in turn could alter the global distribution of the symbiont and alter its global contributions to biological nitrogen fixation.

DNA sequencing in the context of a long time series project, as we report here, is a particularly valuable method for studying microbial interactions and changes therein. In our time series, the 16S/ 18S small subunit of ribosomal RNA (SSU rRNA) genes of the entire microbial community are processed into ASVs using “universal” primers that capture all three domains of life [29–31]. Because we did not design the study with a particular set of organisms in mind, we can retrospectively examine these data for potential biological interactions of any microbes. Each set of DNA sequences from a community provides a single snapshot into its structure and function at the time of sampling. Thus, long time-series projects, in which communities are probed on a regular basis over years or decades, are particularly powerful: they allow researchers to assemble a “movie” of what entire microbial communities are actually doing over time [32], including looking retrospectively at events that may have occurred only rarely. In this study, we sought to characterize the abiotic niche of UCYN-A, its potential predators, and its host specificity using 16S/ 18S amplicon sequence variants over the course of a decade-long time series in coastal, temperate waters.

## Materials and Methods

### Data collection

Seawater was collected monthly from 5m (surface) and deep chlorophyll maxima (DCM) from 2000-2018 at the San Pedro Ocean Time-series (33.55°N, 118.4°W; Figure 1). Samples were collected and processed into 16S/ 18S Amplicon Sequence Variants (ASVs, which differ by as little as one base pair) as described in Yeh and Fuhrman ([33], also see supplemental methods for more details). For the analyses reported here, ASVs corresponding to chloroplasts and multicellular metazoans were removed.

**Figure 1.**
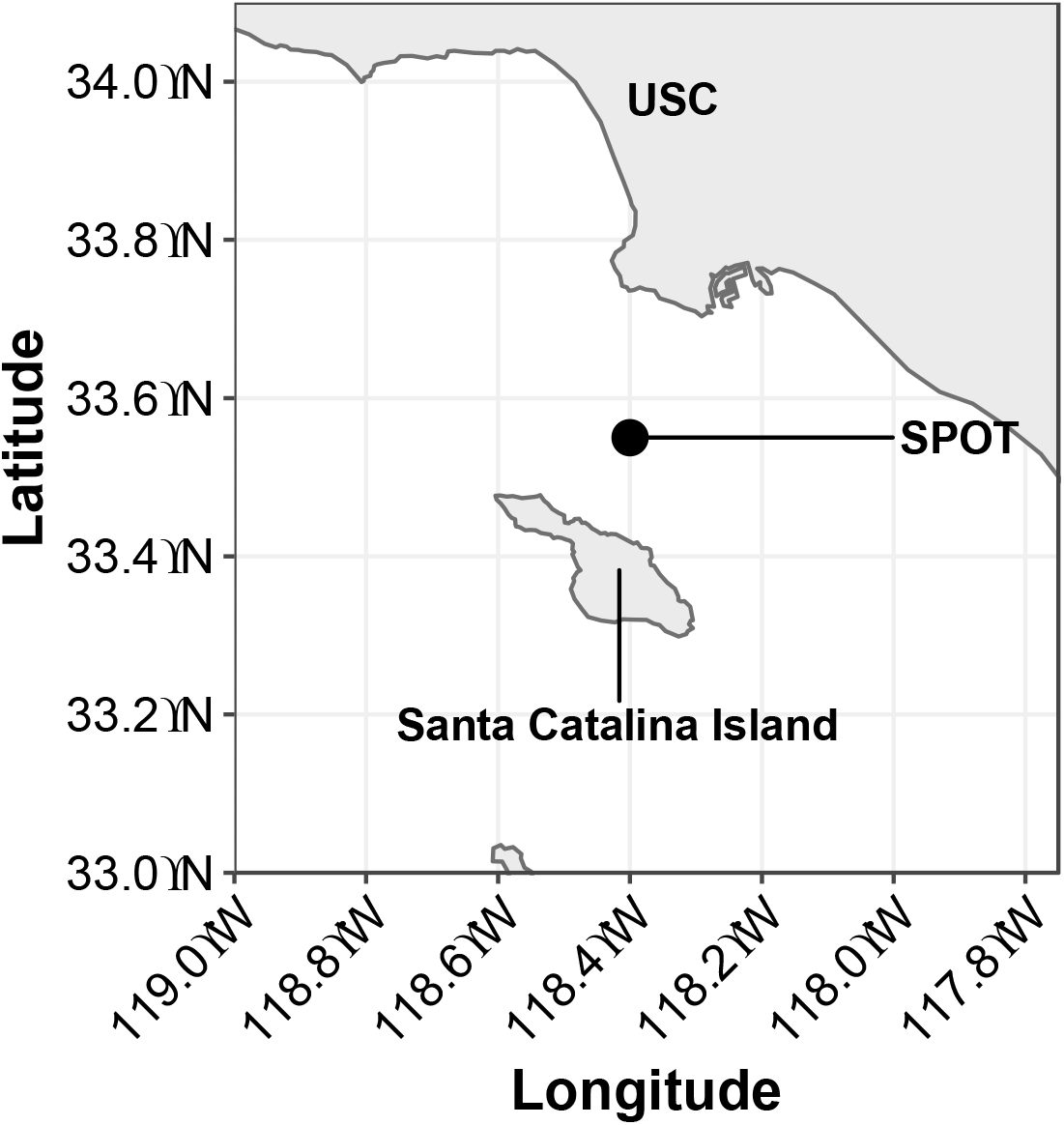
Sampling site location for the San Pedro Ocean Time-series (SPOT). SPOT is ~16km from the Port of Los Angeles (33.55°N, 118.4°W). USC, University of Southern California. Sample collection is described in detail in Yeh et al. [33].

To investigate host specificity, relative abundances of UCYN-A ASVs and their known hosts were normalized to a common denominator and compared (see supplemental methods).

Upwelling intensity at 33°N, measured by improved indices for the U.S. West Coast, the Biologically Efficient Upwelling Transport Index (BEUTI) and Coastal Upwelling Transport Index (CUTI) [34], were downloaded from the National Ocean and Atmospheric Administration (NOAA’s) Pacific Fisheries Environmental Laboratory (https://oceanview.pfeg.noaa.gov/products/upwelling/cutibeuti). Components of the Bakun Index for upwelling at the nearest available point (33.5°N, −118.5°W) were also downloaded from NOAA (https://coastwatch.pfeg.noaa.gov/erddap/griddap/erdlasFnWPr.html). Multivariate ENSO Index (MEI) data, indicative of El Niño (positive MEI)/ La Niña (negative MEI), were obtained from the NOAA Physical Sciences Laboratory (https://psl.noaa.gov/enso/). As recommended, each bimonthly sliding window was used to represent the latter of the two months. Bacterial production was measured by incorporation of tritiated leucine (e.g. [35]). Inorganic nitrogen ([NO_2_^-^+NO_3_^-^]) and phosphate ([PO_4_^3-^]) concentrations at the time of sampling were measured via a LACHAT spectrophotometer QuickChem 8500 Series 2 at the Marine Science Institute at the University of California, Santa Barbara (concentration range 0.2-300μM for nitrogen, 0.1-200μM for phosphate).

### Phylogenetic tree

QIIME2 classified 16S ASVs as UCYN-A with >99.9% confidence, while seven 18S ASVs were classified as *Braarudospharea* with >93% confidence. These ASVs, along with published 16S sequences of UCYN-A and 18S sequences of *Braarudospharea* were assembled into a phylogenetic tree for host and symbiont. Trees included all UCYN-A and *Braarudospharea* sequences publicly available as of January 2022. Sequences were aligned via maaft with default settings [36]. The alignment was trimmed via trimal, also with default settings [37]. Trees were constructed via RAxML (Randomized Accelerated Maximum Likelihood) with rapid bootstrap analysis and 100 bootstraps [38], and visualized via the interactive Tree of Life (iTOL; [39]).

### Relationship with abiotic factors

Logistic regression was used to evaluate effects of each environmental parameter on UCYN-A symbiont and host presence at SPOT. This process was repeated for a *Lepidodinium* ASV. The mean and standard error of each parameter were plotted on days that each organism was present/ absent (defined as >0.01% of the 16S community) via ggplot 2 [40]. Statistical significance of these differences was corrected for multiple testing via Benjamini-Hochberg correction.

For all other analyses, relative abundance data were prepared as follows. Five samples from the 5m depth and two samples from the DCM were excluded because they contained too few sequences to capture the diversity of 18S ASVs at SPOT (Figure S1). Abundance data from missing dates were linearly interpolated via na.approx() from the R package zoo [41]. To avoid common problems with compositional data [42, 43], relative abundances were CLR transformed with the mclr() function from the SPRING package [44].

Spearman’s rank-order correlation was used to evaluate the monotonic relationship between environmental variables and CLR-transformed host/ symbiont relative abundances. Spearman’s correlation was performed using the rcor() function with type=”spearman” from the Hmisc package in R.

Additional analyses are described in Supplementary Methods.

### Network analyses

Co-occurrence networks were generated on interpolated, CLR-transformed data with extended local similarity analysis (eLSA; [45, 46]). Networks constructed using non-CLR transformed data missed several correlations detectable from transformed data (Figure S2). This study reports on associations between UCYN-A and 18S taxa at 5m from March 2008–July 2018. CLR-transformed UCYN-A1 relative abundances from the smaller size fraction were included on all dates. On several dates that UCYN-A1 and UCYN-A2 relative abundances peaked in the larger size fraction prokaryotic community, up to 100 of the most abundant eukaryotic taxa were selected for inclusion in network analyses. eLSAs were run with 1000 permutations and default normalization off. Q values were calculated from P-values using the qvalues() package (e.g. [47]). Correlations at the 5m depth that were highly statistically significant via both Pearson’s correlation and Spearman’s correlation (all P values < 0.005 and all Q values <0.01) were visualized using Cytoscape v3.5 [48]. Relative abundances of the 18S taxa included in these networks were visualized via Krona plots.

## Results

### Phylogenetic tree

Six 16S ASVs were classified as UCYN-A with >99.99% confidence according to QIIME2. ASV names here refer to the alphabetical order of the identifiers generated by QIIME2 (e.g. UCYN-A ASV1 is first alphabetically in a list of six UCYN-A identifiers). SPOT UCYN-A ASV1 matched the published genome of UCYN-A Clade 1 perfectly (Figure 2A; [49]), and will be referred to as UCYN-A1. SPOT UCYN-A ASV5 matched the published genome of UCYN-A Clade 2 (Figure 2A; [14]). In addition, UCYN-A ASV5 never appeared in the smaller size fraction of filters (0.22-1 μm), which is consistent with the reported larger diameter of UCYN-A2 (>1μm) [50]. This ASV will be referred to as UCYN-A2.

**Figure 2.**
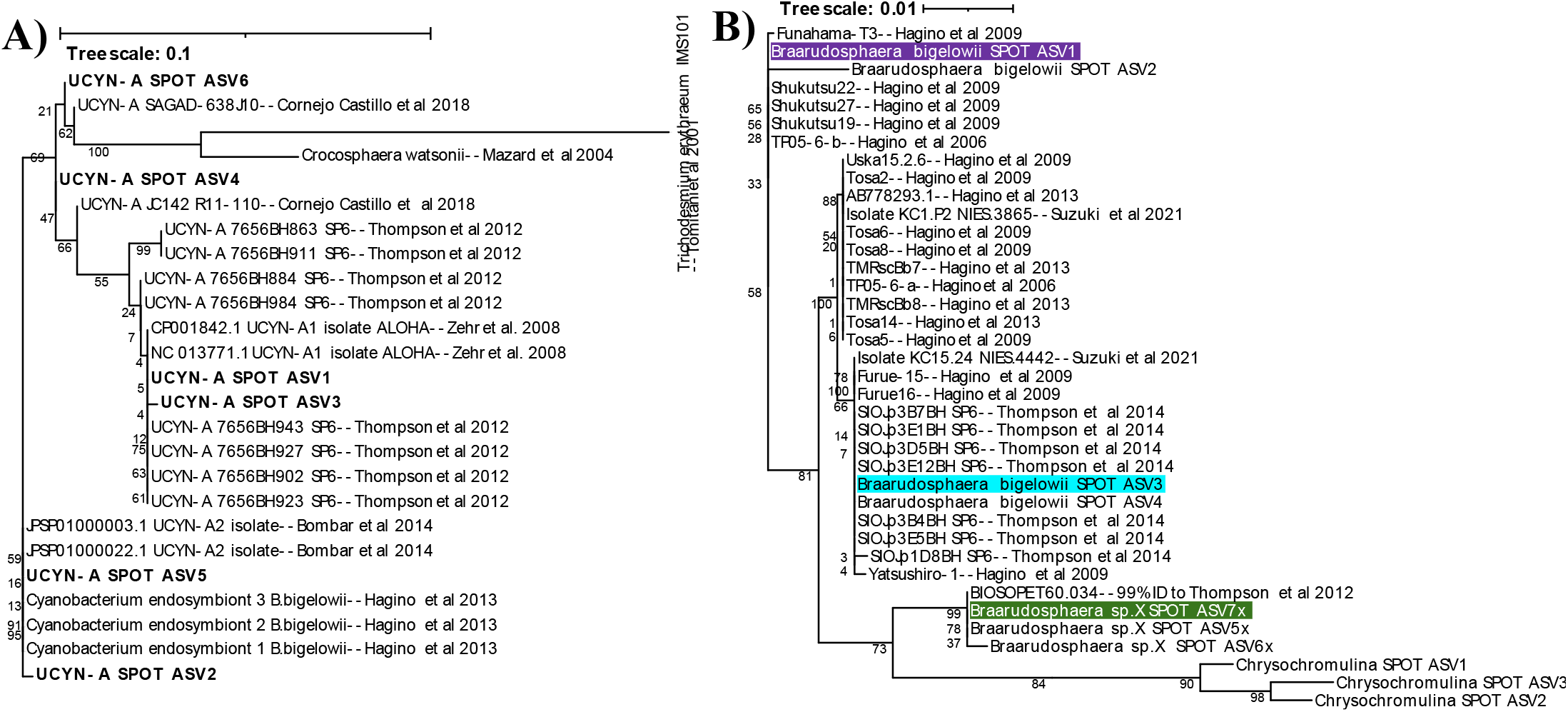
UCYN-A 16S sequences (A) and *Braarudosphearea* 18S sequences (B) from SPOT are phylogenetically identical to or very close to the accepted sequences for UCYN-A1, UCYN-A2, and hosts shown to associate with these symbionts. Bolded names indicate UCYN-A sequences from SPOT; yellow highlighted names indicate 16S sequences from the published genomes of symbionts. *Braarudospharea* ASVs that associate with UCYN-A are bolded and highlighted in green, purple, and blue. Trees were generated via RAxML with 100 bootstraps; numbers indicate bootstrap values.

QIIME2 classified seven of the 18S ASVs found at SPOT as *Braarudosphaera*, the genus that contains previously established hosts of UCYN-A symbionts, with >93% confidence. Four of these seven ASVs were classified as *B. bigelowii*, the putative host of UCYN-A2, and *B. bigelowii* ASV1, ASV3, and ASV4 closely matched strains of *B. bigelowii* shown to associate with UCYN-A2 (Figure 2B; [12]). The other three *Braarudospharea* ASVs did not match a formally named species, and are referred to as 5x, 6x, and 7x. Of these, *Braarudosopharea* ASV7x closely matched isolate BIOSOPE T60, the established host of the published UCYN-A1 genome [8].

### Relationship with abiotic factors

Regional coastal upwelling, measured by BEUTI, was significantly weaker on dates that UCYN-A1 was present (Figure 3, Table S1). In addition, chlorophyll-a concentrations and bacterial production were lower when UCYN-A1 was present, while sea surface temperature (SST) and MEI were higher (Figure 3). Ekman transport moved surface water North and East during months UCYN-A1 was present at SPOT (Table S1). UCYN-A1 relative abundances correlated positively with MEI and SST, and negatively with bacterial production, upwelling, and chlorophyll-a (Figure 4). Higher than average relative abundances of UCYN-A1 coincided with low upwelling indices, positive MEI, and SST >19°C (Figure S4). On dates UCYN-A2 was present, the BEUTI index was slightly lower, indicating weaker upwelling was favorable to UCYN-A2 presence (Figure 3). Relative abundances of UCYN-A2 showed a weak positive correlation with SST and a weak negative correlation with BEUTI (Figure 4), but no significant correlation to chlorophyll concentrations, bacterial production, SST, and MEI (Table S1). Neither nitrogen nor phosphorus concentrations differed significantly on dates that UCYN-A ASVs were present vs. absent (Table S1).

**Figure 3.**
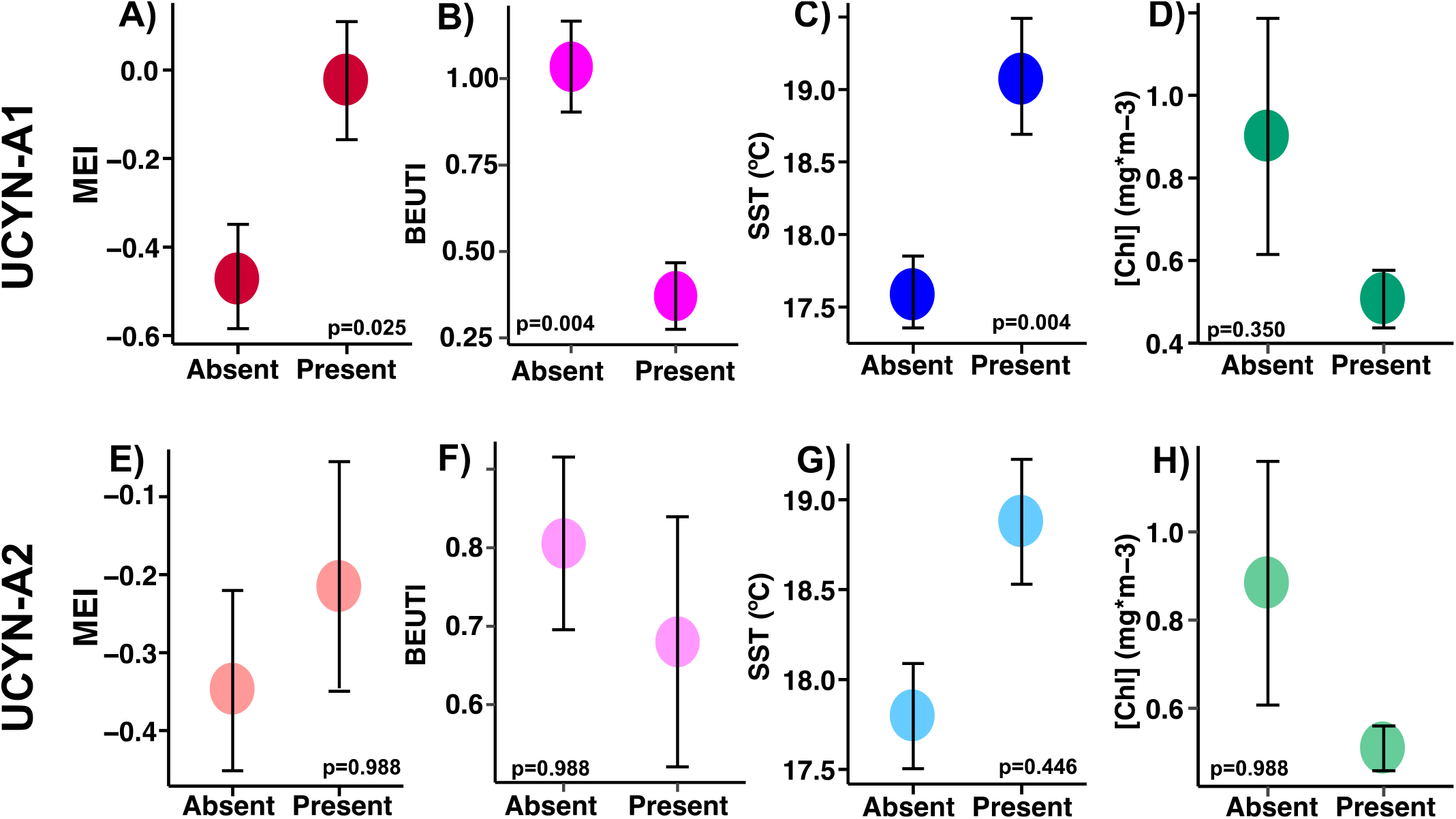
UCYN-A1 is present at SPOT under El Niño conditions, with relaxed upwelling, higher temperatures, and lower biomass (A-D); these patterns are similar but weaker for UCYN-A2 (note larger p-values and error bars). Differences in the mean of each value when ASVs were present/ absent were evaluated via logistic regression. P-values reported were corrected for multiple testing via Benjamini-Hochberg correction. Error bars represent standard error. Higher Multivariate ENSO Index (MEI) indicates El Niño conditions (more stratified); higher Biologically Effective Upwelling Transport Index (BEUTI) indicates stronger upwelling.

**Figure 4.**
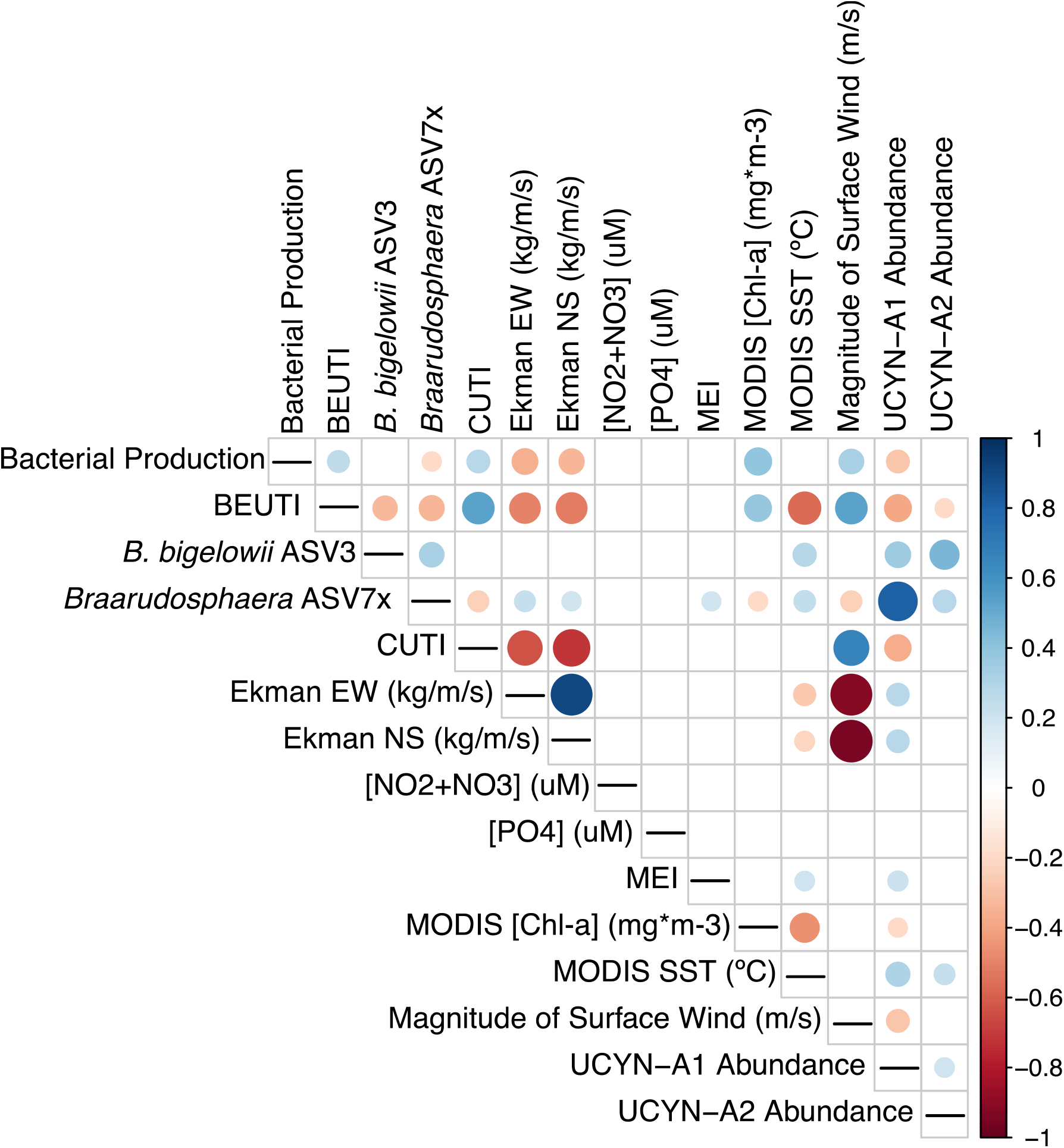
UCYN-A1 relative abundances correlate positively with host organisms, UCYN-A2 host/symbionts, MEI, and SST, and negatively with upwelling indices (BEUTI and CUTI) and indirect indicators of upwelling, including bacterial production, East-West Ekman transport, and chlorophyll concentrations. UCYN-A symbiont/host ASVs were CLR-transformed and correlated with environmental parameters via Spearman’s rank-order correlation. Dot size indicates the strength of correlation, while dot color represents positive or negative associations. BEUTI, Biologically Effective Upwelling Transport Index; CUTI, coastal upwelling transport index; EW, East-West; NS, North-South; MEI, Multivariate ENSO Index.

### Co-occurrence with non-host 18S taxa

With high statistical significance (P<0.005 and Q<0.01 by Local Similarity, Pearson’s and Spearman’s correlation), UCYN-A co-occurred with numerous 18S ASVs, such as prymnesiophytes and Dinoflagellates, notably including parasitic Syndiniales and an ASV from the genus *Lepidodinium*. UCYN-A1 and UCYN-A2 shared several of these taxa in common (Figure 5, Figure S6, Table S2). The *Lepidodinium* ASV was present at SPOT under environmental conditions similar to those found when UCYN-A ASVs were present (Table S1).

**Figure 5.**
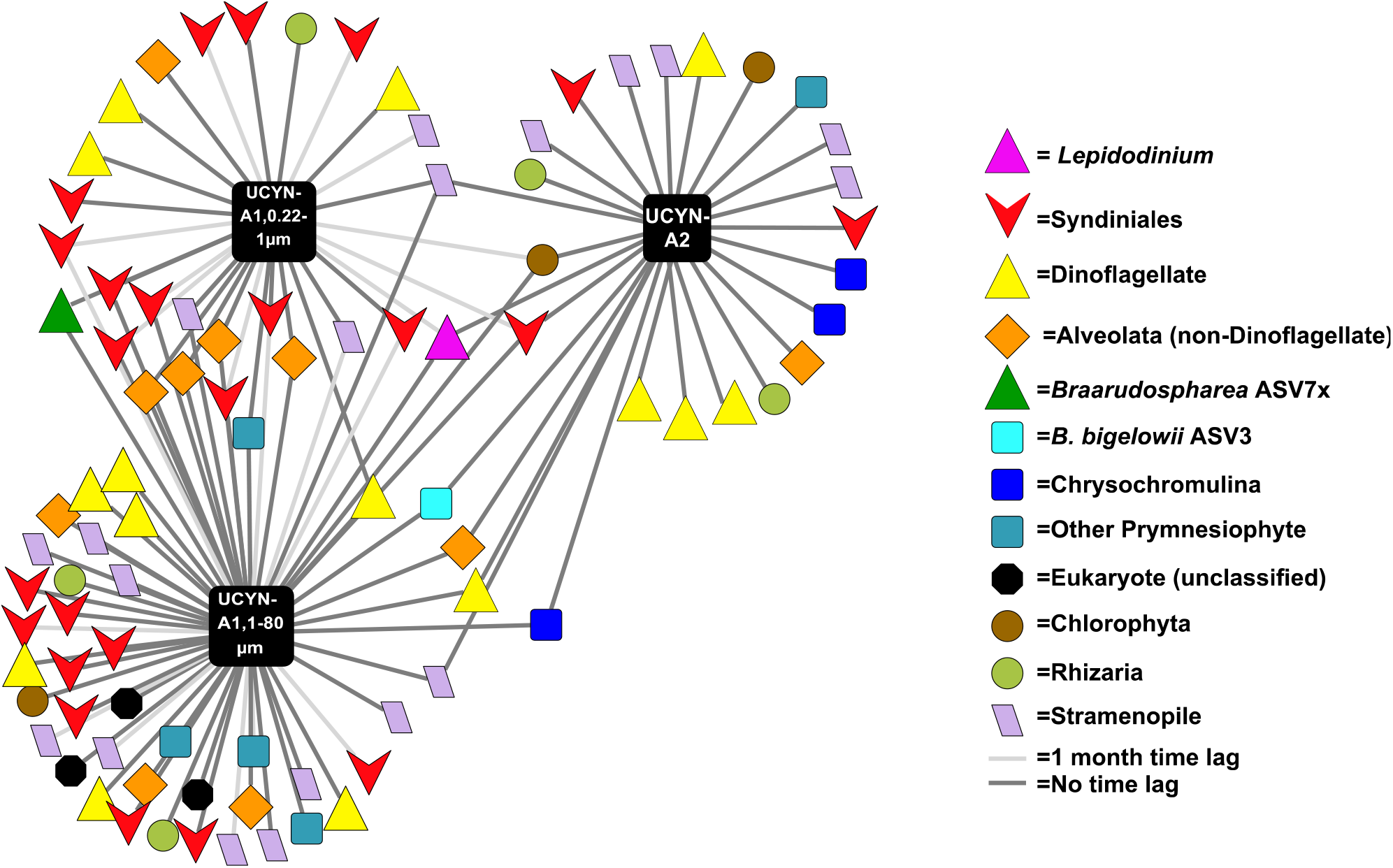
UCYN-A co-occurs with a variety of 18S taxa at the SPOT surface, notably including *Lepidodinium*, hypothesized to be a predator (pink triangle, center). Networks were generated via eLSA and visualized in Cytoscape 3.5. Each node represents one ASV; only ASVs that co-occurred with P<0.005 and Q<0.01 by both Pearson’s and Spearman’s correlation are shown using easily recognizable names. QIIME-generated hashes of each node are presented in Table S2; 16S/ 18S sequences of each node, including UCYN-A symbionts, are publicly available (see Data Availability Statement).

### Host specificity of UCYN-A ASVs

UCYN-A2 co-occurred with *B. bigelowii* ASV3 on ~60% of the dates it was present at the SPOT surface (Figure 6A-C, S7). On average, the ratio of UCYN-A2 16S: *B. bigelowii* ASV3 18S was about 2:1 at the SPOT surface (Figure 6C). Another ASV of *B. bigelowii*, ASV1, peaked in relative abundance on dates where UCYN-A2 was present, but *B. bigelowii* ASV3 was absent (Figure 6D). At the DCM, *Braarudospharea* ASV3 was present on ~60% of the dates UCYN-A2 was present, but the ratio of their 16S: 18S genes of these organisms was about 1:1 on average (Figure S8).

**Figure 6.**
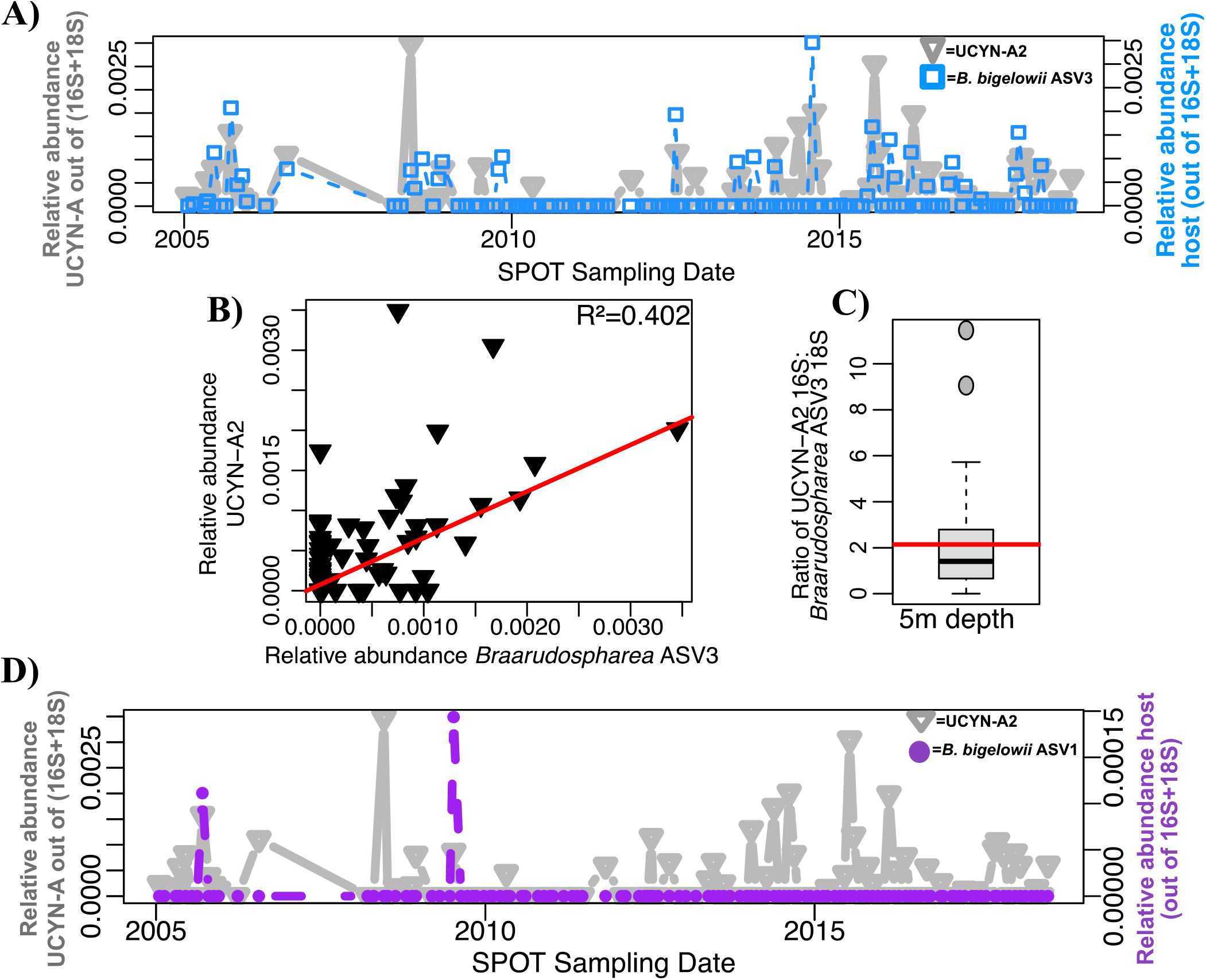
UCYN-A2 is less tightly coupled to its prymnesiophyte host than its Clade 1 counterpart. A) UCYN-A2 often, but not always, co-occurs with its most common host at 5m depth across the SPOT time series. B) UCYN-A2 relative abundance correlates with relative abundance of its most common host more weakly than the Clade 1 symbiosis. C) The ratio of 16S: 18S genes of these organisms is lower than expected. Boxplot values indicate the median and interquartile range values of this ratio; the red line indicates the average (2.139). 29 sampling dates, on which host and symbiont are present, are included. D) UCYN-A2 occasionally co-occurs with *Braarudospharea* ASV1. The relative abundances of 16S and 18S ASVs were normalized as described in Supplementary Methods to allow for direct comparison.

The relative abundance of UCYN-A1 closely mirrored that of *Braarudosphaera* ASV7x, the closest match for the known host of UCYN-A1 (BIOSOPE T60; Figure 2B), at the SPOT surface (Figure 7A-C, S7). The 16S gene of UCYN-A1 and 18S gene of *Braarudosphaera* ASV7x consistently co-occurred in nearly a 2:1 ratio at 5m depth (Figure 7C). In addition, UCYN-A1 co-occurred with *Braarudosphaera* ASV7x in eLSA networks with high statistical significance at the SPOT surface (Figure 5). These organisms mirror each other less strongly at the DCM, where their ratio of 16S: 18S genes is also approximately 2:1 (Figure S9).

**Figure 7.**
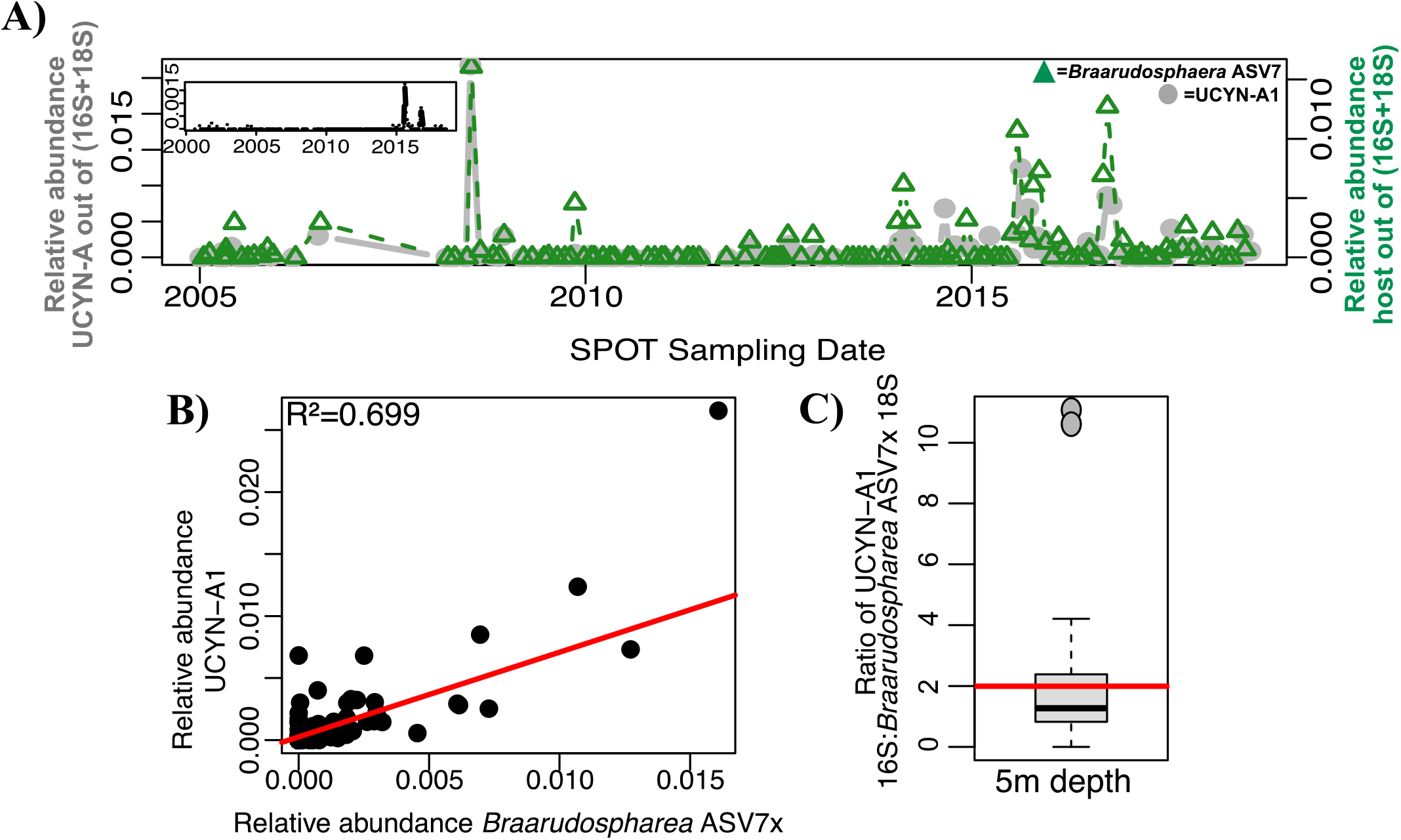
UCYN-A1 and its prymnesiophyte host are tightly coupled throughout the SPOT dataset. A) UCYN-A1 consistently co-occurs with its putative host at 5m depth across the SPOT time series. Panel inset indicates the relative abundance of UCYN-A1 in the smaller size fraction. B) UCYN-A1 relative abundance strongly correlates with its putative host relative abundance. C) The ratio of 16S: 18S genes of these organisms is, on average, as expected. Boxplot values indicate the median and IQR values of this ratio; the red line indicates the average (1.991). One outlier (ratio >50) was excluded. 43 sampling dates are included. As with UCYN-A2, the relative abundances of 16S and 18S ASVs were normalized to allow for direct comparison.

### Spatial-temporal distributions of UCYN-A ASVs

UCYN-A ASVs were primarily found in the larger size fraction in the euphotic zone and were also detected once at 890m at about 0.5% of the 16S community (Figure S3, S10). UCYN-A1 was found on 44% of sampling dates, while UCYN-A2 was present at the SPOT surface on 38.4% of all sampling dates (Figure S3, 6A, 7A). Relative abundances of all *Braarudosphaera* ASVs were highest in the euphotic zone, and they appeared at depths greater than the DCM even more rarely than UCYN-A.

## Discussion

### Relationship with abiotic factors

Currents flowing from the tropical Pacific to the North and East likely brought the UCYN-A1 symbiosis, the “open ocean” ecotype, into our study system. These currents are particularly strong during El Niño conditions and less so during regional upwelling. Positive Multivariate ENSO Index (MEI) values indicate El Niño conditions, in which warm, tropical waters flow West to East across the equator, altering climate patterns throughout the Pacific Ocean basin and weakening upwelling (e.g. [51, 52]). UCYN-A1 presence and relative abundance at our study site are positively correlated with higher MEI, i.e. El Niño conditions (Figure 3, Table S1, Figure 4, Figure S4). Our data show a sudden increase in the relative abundances of UCYN-A1 and its host following the 2015-2016 El Niño event, which were sustained in the subsequent years (Figure 7A). Similarly, other time-series data show offshore ASVs were advected into the Southern California Bight after the same El Niño event [53], including warm ecotypes of Prochlorococcus at our study site (Yeh et al., in prep). Prior to this event, e.g. in 2010-2011, the UCYN-A1 symbiosis was infrequently detected at our sampling site and was similarly absent from other locations in the Southern California Current System (SCCS) during overlapping periods of sampling [13, 22].

In addition, multiple lines of evidence suggest that UCYN-A1 presence and relative abundance are negatively influenced by seasonal upwelling. In this annual, coastal event, Ekman transport displaces surface waters West and South from SPOT, and nutrient-rich waters rise from the deep to replace them, stimulating increased bacterial chlorophyll concentrations and productivity (e.g. [34, 54]). Upwelling is here measured by the Biologically Effective Upwelling Transport Index (BEUTI) and Coastal Upwelling Transport Index (CUTI) [34], with higher BEUTI and CUTI values indicating stronger upwelling. Lower BEUTI and CUTI are correlated with UCYN-A1 presence (Figure 3, Table S1), while sparse binomial regression indicated that the CUTI index negatively predicted UCYN-A1 presence at SPOT (see Supplemental Information). Indirect indicators of seasonal upwelling, including Ekman transport to the South and West, lower SST, higher bacterial production rates and [Chl-a], were also negatively correlated with UCYN-A1 presence and abundance (Figure 3, Table S1, Figure 4, Figure S4). Similarly, UCYN-A and hosts regularly appeared in Monterey Bay, North of our study site, in August-November, after regional upwelling had ceased [22]. Previous studies have also related UCYN-A1 abundances to temperature: although UCYN-A distribution may not be directly influenced by temperature [9, 19], warmer water conditions are favorable to increased UCYN-A1 relative abundance [21, 22]. In tandem, these observations indicate that the “open ocean” ecotype of the UCYN-A symbiosis is advected into our study system by warm, tropical waters during El Niño and periods of relaxed upwelling, which foster lower biomass and coincide with higher temperatures, providing ideal conditions for the open-ocean ecotype to proliferate.

UCYN-A1 relative abundances and nitrogen fixation rates were observed to increase days after upwelling in nearshore samples from the Scripps Pier and the Alaskan Beaufort shelf [21, 55]. The daily sampling resolution of these cruises or their proximity to shore may explain discrepancies with our study. Upwelling may stimulate UCYN-A seed populations at a daily timescale, perhaps due to community production of a micronutrient not measured here that is essential to the UCYN-A1 symbiont or host [21]. Due to the monthly resolution of our sample collection regime, SPOT data may not capture these short-term dynamics. Instead, we observe that upwelling hinders UCYN-A proliferation on monthly timescales. Notably, both previous studies collected samples solely in 2017, soon after the 2015-2016 El Niño event. This could have brought UCYN-A seed populations to the Scripps Pier study, which was concurrent with an influx of warm, low-density waters from the tropics. Our study shows that over longer time scales, ocean currents including this El Niño event and relaxed upwelling created warmer, low biomass conditions which were conducive to UCYN-A1.

Differences between UCYN-A1 and UCYN-A2 symbioses with respect to their abiotic niches are consistent with the hypothesis partitioning the two clades into coastal and open-ocean ecotypes (e.g. [14]). UCYN-A2 presence was not strongly affected by seasonal upwelling or El Niño (Figure 3), and its relative abundance was only correlated with two abiotic factors: BEUTI index (negative) and SST (positive) (Figure 3, Table S1, Figure S4).

Given the paradigm presented here, climate change will likely have a strong but mixed effect on UCYN-A1 proliferation and activity at SPOT. Intense El Niño events are expected to become more frequent with climate change [57], advecting warm, tropical waters, along with UCYN-A1, into Southern California more often. Simultaneously, upwelling events along the California Current System are expected to intensify, as warming on land increases air pressure differences between land and sea, strengthening coastal winds and therefore upwelling [54, 58, 59]. Notably, the degree to which upwelling may intensify is unclear [60], partially due to mitigative effects of El Niño events, which weaken upwelling [52]. Although upwelling deters UCYN-A1 proliferation at SPOT, the symbiosis was able to recover from upwelling events following the 2015-2016 El Niño event (Figure 7). Future populations of UCYN-A1 may adhere to this trend, such that the symbiosis reaches greater abundances following future El Niño events. Because UCYN-A symbioses comprise up to 95% of the nitrogen fixing population in Southern California [21], this could substantially increase inputs of fixed nitrogen to this region.

### Co-occurrence with non-host 18S taxa

Based on co-occurrence data with high statistical significance (Figure 5, S6, Table S2), we hypothesize a predator-prey relationship between the UCYN-A symbiosis and *Lepidodinium*, which in turn may be parasitized by Syndiniales (Figure S6). The dinoflagellate genus *Lepidodinium* has been shown to prey on the tropical nitrogen fixer *Crocosphearea watsonii* through Lotka-Volterra modeling of environmental data [27], and doubles its grazing rates at night, when *C. watsonii* organism fixes nitrogen [26]. Furthermore, *Lepidodinium* has been hypothesized as a grazer of UCYN-A symbioses, as the diameter of UCYN-A symbioses (1-3μm for UCYN-A1 symbioses and 4-10μm for UCYN-A2 symbioses) are similar to that of *C. watsonii* (3.9μm for *C. watsonii* [26]). Its appearance in our dataset supports this hypothesis, suggesting that this genus could prey on UCYN-A and other nitrogen fixers. The order Syndiniales is primarily known to parasitize dinoflagellates (e.g. [61, 62]), and its members can be generalists or specialists [63]. Syndiniales ASVs most likely co-occur with UCYN-A indirectly, due to the presence of their dinoflagellate hosts (see Figure S6). However, it is possible that Syndiniales are parasitizing *Braarudospharea* directly, as some Syndiniales are able to parasitize non-dinoflagellate hosts from genetically distinct phyla (e.g. [64]). Our co-occurrence data and others show statistically significant co-occurrence links between haptophytes and Syndiniales (Figure 5, Figure S6, Table S2 [65]), and are consistent with the possibility of parasitism on a haptophyte.

Many of the associations discussed here have been reported previously. The paper that announced *Braarudosphaera* as the putative host of UCYN-A1 used a UCYN-A *nifH* probe and FACS (Fluorescent-Activated Cell Sorting) to show that UCYN-A physically associates with many families, including Dinophyceae, the family that contains Dinoflagellates [8]. Using 18S primers, Krupke et al. found that UCYN-A co-occurs with Rhizaria, Dinoflagellates, including *Lepidodinium* and Syndiniales, and prymnesiophytes including *Chrysochromulina*, which is taxonomically very close to (and in some cases may overlap with) *Braarudospharea* [65]. One recent study proposed that the prymnesiophyte called *Chrysochromulina parkaea* is an uncalcified life stage of *B. bigelowii* (99.87% identical 18S sequence to a *Braarudosphaera* sequence) and is also able to associate with UCYN-A [67].

It should be noted that co-occurrence of taxa does not imply a biological association [68]. It is possible that UCYN-A and *Braarudosphaera* do not interact with any of the taxa reported here, but happen to co-occur due to a mutually favorable environmental niche. Notably, the *Lepidodinium* ASV is present at SPOT under similar conditions as UCYN-A1 and UCYN-A2 (Table S1). However, few of these parameters differed significantly on dates *Lepidodinium* was present vs. absent, and it is unclear if they influence the organism’s relative abundance. Additional studies should be performed to elucidate true interactions. For example, CARD FISH with labelled probes for *Lepidodinium* and for UCYN-A could reveal physical evidence of predation. To tease apart other potential interactions, multi-stressor grow-out experiments could be conducted, in which temperature and availability of key nutrients are each varied along a gradient (e.g. [69]).

### Host specificity of UCYN-A ASVs

At the 16S ASV level, UCYN-A2 is not as tightly coupled with its prymnesiophyte partner as is UCYN-A1. UCYN-A2 frequently appeared at the SPOT surface in excess of its presumed host, based on ratios of their 16S: 18S genes. The UCYN-A2 host is thought to have 30-40 copies of the 18S rRNA gene [14], and 7-10 symbionts [16], such that the ratio of UCYN-A 16S rRNA genes: 18S *Braarudosphaera* host genes should be about 0.66 at most. Due to differences in copy number, as well as different stages of cell division for symbiont or host, we do not expect a constant ratio or a linear relationship between host 18S and symbiont 16S genes. However, the average ratio of sequences was much higher than expected (average=2.729), and their genes showed poor correlation (Figure 6B-C).

We also show strong circumstantial evidence that UCYN-A2 can partner with another prymnesiophyte host, *B. bigelowii* ASV1. On two days in particular, UCYN-A2 peaked in relative abundance when the abundance of its presumed host, *B. bigelowii* ASV3, was lower than expected. On both of these days, and on no other days, *B. bigelowii* ASV1 also peaked in relative abundance (Figure 6C). Both this ASV of *B. bigelowii* and ASV3, the predominant host, are closely related to genotypes of *Braarudosphaera* found to associate with UCYN-A2 in the coastal waters of Japan. Hagino et al. used full length 18S sequencing, CARD FISH, and electron microscopy to observe that multiple cryptic species of *B. bigelowii* from disparate habitats in the Japanese islands possessed UCYN-A symbionts [12, 17]. Notably, these two *Braarudosphaera* ASVs cluster with different pseudo-species: *B. bigelowii* ASV1 clusters with Intermediate form 1A, Genotype I, while *B. bigelowii* ASV3 co-occurs with Large form, Genotype IV [17] (Figure 2B). The UCYN-A2 ASV we observed may have different partners in different environmental conditions. We may have detected a “local” pair that is usually rare and became abundant on certain dates or might have been temporarily brought to SPOT from a distant location on the dates alternate hosts were observed. It is also possible that these two hosts have different symbionts that are identical at the 16S ASV level.

Regardless, UCYN-A2 and its host are not present at SPOT under the same environmental conditions: UCYN-A2 was present at the SPOT surface more frequently than its reported host (38.4% vs. 28% of sampling dates, Figure 6A). Compared to its host, the UCYN-A2 symbiont was present on days with increased concentrations of phosphate and increased upwelling (currents flowing to the South and West), although these differences are not statistically significant (Table S1). Culture-based studies support the idea of weak coupling between UCYN-A2 symbiont and host, noting that the UCYN-A2 host appears able to dissociate from its N_2_-fixing symbiont and rely on predation as a nitrogen source [67].

In contrast, UCYN-A1 appeared tightly coupled to its established *Braarudosphaera* host. On >75% of days UCYN-A1 or *Braarudosphaera* ASV7x were present at SPOT, the two organisms were present in an average ratio of 2.176, close to the expected 2:1 ratio of UCYN-A1 16S: *Braarudosphaera* 18S rRNA genes (UCYN-A1 has two 16S rRNA genes and its host is thought to have one copy of the 18S rRNA gene based on its small biomass [14, 16]). Each organism peaked in relative abundance only on days that its partner was also present. Conversely, on dates that one half of the pair was present but the other absent, the organism was only present in low abundances. These two organisms were present under similar environmental conditions (Table S1), and co-occurred with high levels of statistical significance (Figure 5). This adds to existing evidence of the tight metabolic partnership between UCYN-A1 and its haptophyte host (e.g. [50, 56]).

Another ASV from UCYN-A Clade 1, ASV6, was also present at the SPOT surface (Figure S3), but did not co-occur with any *Braarudosphaera* ASVs at any level of statistical significance. Its relative abundance peaked on dates that the known UCYN-A hosts were absent (Figure S7, S13). Other studies have reported free-living UCYN-A1 symbionts in open-ocean regions but attributed this phenomenon to host-symbiont dissociation during sample collection [19]. However, these studies used CARD-FISH and clustered 16S sequences into operational taxonomic units (OTUs) rather than ASVs, methods that miss the high-resolution differences between 16S ASVs from the same clade of organisms. This further illustrates the utility of high-resolution 16S ASVs (e.g. [24]).

### Spatial-temporal distributions of UCYN-A ASVs

As expected, UCYN-A ASVs were primarily found at 5m and the DCM throughout the time series (Figure S3, S10), consistent with reports that its host is photosynthetic [8]. However, UCYN-A1 was found as deep as 890m depth on least one instance (Figure S10), and at around the same time prior studies noted the UCYN-A *nifH* gene at the bottom of the San Pedro channel [23]. This strengthens existing evidence of the host-symbiont organisms’ capacity to export calcareous carbonate and fixed nitrogen from the euphotic zone [8], although this study does not address whether nitrogen fixation occurs at the depths. *Braarudosphaera* ASVs were found almost exclusively in the euphotic zone; the limited depth range of this genus is unique amongst Prymnesiophytes, most of which can proliferate at depths deeper than the DCM [9, 18].

Both the coastal and open ocean ecotypes of UCYN-A were present in our SPOT dataset, reflective of the fact that SPOT is an open ocean sampling site but is close enough to shore to be influenced by coastal dynamics [70]. UCYN-A1 appeared in 44% of all large size fraction samples collected from 5m depth at SPOT (1-80μm), and often in relative abundances as high as 2.5% of the whole community (Figure S3, 7). On several occasions, UCYN-A1 was found in very low relative abundances in the smaller size class of organisms (0.22-1 μm) (Figure 7A, inset). The UCYN-A1 symbiont is known to dissociate from its host, most likely due to the gentle pressures of sample filtration, which explains why it is found in multiple size fractions [11].

UCYN-A2 appeared in almost as many SPOT surface samples as UCYN-A1 (38.4%), but in lower relative abundances, at most 0.3% of the community (Figure S3, 6). This ASV was never found in the smaller size fraction, which further indicates that the ASV is from UCYN-A2: UCYN-A Clade 2 organisms are reported to have a diameter >1μm [50] and should consistently remain in the larger size fraction (1-80μm). The UCYN-A2 symbiosis was outcompeted by its Clade 1 counterpart at our study location: both were present on ~40% of sampling dates (Figure S3, 6A, 7A) and co-occurred with one another (Figure 4, Figure S7), but UCYN-A1 was present at almost an order of magnitude higher relative abundance than UCYN-A2 (at most 2.5% of the community vs. 0.3%; Figure 7A, 6A). This may be because the “coastal ecotype” is accustomed to living in environments with more biomass than is typically present at SPOT, and the study location is more comparable to environments preferred by the “open ocean ecotype.”

### Conclusions

This paper reports trends in the spatio-temporal dynamics of the symbiotic diazotroph UCYN-A, its haptophyte hosts, and other associated taxa over ten years off the California Coast. We present important differences between two highly studied clades of UCYN-A with regards to host specificity, co-occurrences with potential predators, and abiotic parameters. Studying these changes is particularly important as climate change continues to alter the distributions of UCYN-A, its hosts, and other associated 18S taxa [2].

## Supporting information

Interactive HTML file with 18S taxa that co-occur with all UCYN-A ASVs

Interactive HTML file with 18S taxa that co-occur with UCYN-A1

Interactive HTML file with 18S taxa that co-occur with UCYN-A2

## Acknowledgements

The SPOT dataset is the collective effort of generations of undergraduate, graduate student, and postdoctoral researchers. We particularly wish to thank Dave Caron, Jed Fuhrman, Troy Gunderson, Diane Kim, and the crew of the R/V Yellowfin for their support. We gratefully acknowledge invaluable conversations with Ana María Cabello, Virginia Edgcomb, Kendra-Turk Kubo, Jon Zehr, and others at the ASLO Ocean Sciences Meetings in 2020 and 2022. This work was supported by NSF OCE 1737409, Gordon and Betty Moore Foundation Marine Microbiology Initiative grant 3779, and Simons Foundation Collaboration on Computational Biogeochemical Modeling of Marine Ecosystems (CBIOMES) grant 549943. Research was conducted on land and waters traditionally stewarded by the Tongva people.

## Author contributions

CFH devised the research questions, led data analysis, data visualization, and interpretation of results, and drafted the manuscript. YCY carried out sample processing and sequencing. YR contributed to data analysis and advised on manuscript writing. JLW advised on statistical analyses. JAF supervised the work and advised on data analysis and manuscript drafting. All authors edited the manuscript.

## Data Availability Statement

Forward and reverse reads from each sample in the San Pedro Ocean Time-series (SPOT) are available at EMBL under accession number PRJEB48162 and PRJEB35673, as described by Yeh and Fuhrman [33]. Scripts necessary to reproduce the analysis are available at https://github.com/jcmcnch/eASV-pipeline-for-515Y-926R [30]. ASV tables generated from these files are available in /OriginalFiles at https://osf.io/6ku49/. Input files for the analyses presented here are available in /ModifiedFiles at the same link. Scripts necessary to reproduce these analyses are available at https://github.com/fletchec99/UCYNA_at_SPOT.

