## Supplementary Material for "Symbiotic diazotrophic UCYN-A strains co-occurred with El Niño, relaxed upwelling, and varied eukaryotes over 10 years off Southern California Bight"

### **Supplementary Information**

#### *Materials and Methods*

DNA SEQUENCING AND PROCESSING: 15-20L of seawater was filtered sequentially through an 80μm mesh (removing mesoplankton) a 1μm glass AE filter (Pall, Port Washington, NY) (collecting the larger 1-80μm size fraction), and finally a 0.2 μm Durapore filter (ED Millipore, Billerica, MA) (collecting the 0.2-1 μm smaller size fraction). DNA was extracted from Durapore filters using the phenol-chloroform method described by Fuhrman et al. [72]. DNA was extracted from AE filters via bead beating, followed by a phenol-chloroform protocol, as described in Lie et al. [73]. The V4-V5 hypervariable region of the 16S and 18S rRNA gene was amplified from these extracts using the universal primers (515Y/926R) as reported by Parada et al. [29], which amplify sequences from both eukaryotic and prokaryotic ribosomal RNA genes [31, 33]. Samples were sequenced on either a HiSeq 2500 in PE250 mode or MiSeq PE300 platform at the USC and UC Davis Genome Core facilities at a target depth of 100,000 sequences/ sample. Per-sample sequence files were submitted to the EMBL database under accession number PRJEB48162 and PRJEB35673. Sequences were processed into Amplicon Sequence Variants (ASVs), which differ by as little as a single base pair, using DADA2 implemented in QIIME2 [74] with scripts available at github.com/jcmcnch/eASV-pipeline-for-515Y-926R. [30]. Prokaryotic and eukaryotic ASVs were taxonomically classified using SILVA 132 in May and July of 2020, respectively.

PRINCIPLE COMPONENT ANALYSES: Principle component analysis (PCA) was used to visualize differences in environmental parameters on dates UCYN-A ASVs were absent (<0.01% of the 16S community) vs. present in relative abundances higher or lower than average. PCA was conducted on abiotic data, which was centered and scaled, using prcomp() in R. Ordinations were plotted using autoplot() from the "ggfortify" package in conjunction with ggplot2 [40].

MODELING EFFECTS OF ENVIRONMENTAL PARAMETERS: Sparse binomial regression was used to resolve which environmental parameters best predicted whether UCYN-A1 and UCYN-A2 would be present at SPOT. Model input data consisted of bacteria production rates, nutrient availability, upwelling indices, and other environmental variables. Data from missing dates were linearly interpolated via na.approx() from the R package zoo [41]. UCYN-A ASVs were considered "present" on dates that they were over 0.01% of the 16S community, and "absent" when their relative abundances were lower than 0.01%. For each ASV, a sparse binomial logistic regression model was constructed via the glmnet package in R [75, 76]. 80% of the data was used as training data for the model, and 20% was used as the test set. Variable selection was performed using lasso regression, and the appropriate lambda was selected using 10-fold cross validation on the training set. F1, sensitivity, specificity, and accuracy of each model were calculated on the test set using the caret package [76].

DATA NORMALIZATION: Our DADA-2 pipeline splits 16S and 18S sequences, generating separate tables of ASVs for prokaryotes and eukaryotes. In order to plot UCYN-A and associated eukaryotes with the same denominator, sequencing data was normalized as follows: Sequences from chloroplasts and metazoans were removed, leaving only SSU sequences from prokaryotes and single-celled eukaryotes in the dataset. Raw sequencing counts of prokaryotic and eukaryotic ASVs were divided by the percent of sequences passing quality control in DADA2. Because HiSeq and MiSeq platforms have been shown to discriminate against the 18S rRNA sequences, favoring the shorter 16S rRNA sequences with a two-fold bias [78], sequence counts from eukaryotic ASVs were then doubled. Normalized sequencing counts of prokaryotic and eukaryotic ASVs were combined and converted to proportions, representing the relative abundances of taxa out of the entire microbial community (16S+18S sequences). This method was developed and successfully tested on mixed mock communities, which contain 16S and 18S rRNA sequences in equal concentrations [78], that were sequenced via HiSeq or MiSeq. Following normalization, communities contained equal proportions of each of the organisms in the sequenced sample, as expected (Figure S11). Code normalizing the 16S/ 18S ASV tables of this QIIME-2 pipeline [30] is available at https://github.com/fletchec99/normalizing_16S_18S_tags.

*Results and Discussion*

PRINCIPLE COMPONENT ANALYSES: Principle component analyses (PCA) indicate that temperature drove variation in the abiotic factors along PC1, which was generally associated with UCYN-A1 and host presence. Higher MEI (Multivariate ENSO Index) was also associated with UCYN-A1 and host presence in PCA. Upwelling indices, as well as indirect indicators of upwelling such as increased nutrient concentration, bacterial production, and chlorophyll concentration, drove variation along PC2 and were associated with UCYN-A1 and host absence. These trends were not as obvious for UCYN-A2 (Figure 4).

MODELING EFFECTS OF ENVIRONMENTAL PARAMETERS: Sparse binomial logistic regression indicated that UCYN-A1 presence was negatively predicted by upwelling and no other variables (coefficient= -0.570, sensitivity=0.667, specificity=0.333, accuracy=0.542, F1=0.645). Models were not able to reliably predict the presence of UCYN-A2 (sensitivity=1.00, specificity=0.00, accuracy= 0.5, F1=0.667).

RELATIONSHIP WITH INORGANIC NUTRIENTS: UCYN-A ASVs were not significantly correlated with inorganic nitrogen and phosphorus concentrations (Table S1, Figure 4). Others have observed UCYN-A abundances and activity have no strong relationships with nitrogen concentrations (e.g. [19, 56, 57]). Due to the well-established link between upwelling and increased inorganic nutrient concentrations (e.g. [34]), the strong influence of upwelling on UCYN-A1 might seem incongruous with the weak influence of inorganic nutrients. It is important to note that upwelling indices are aggregated across latitude on a monthly basis, whereas the inorganic nutrients were measured at SPOT on the day of sampling. Upwelling in Southern California is generally coastal, and it is likely that coastal phytoplankton close to the sites of upwelling consumed the upwelled nitrogen and phosphate, before these compounds could reach our study site, ~16km from the coast (Figure 1).

*Figures*

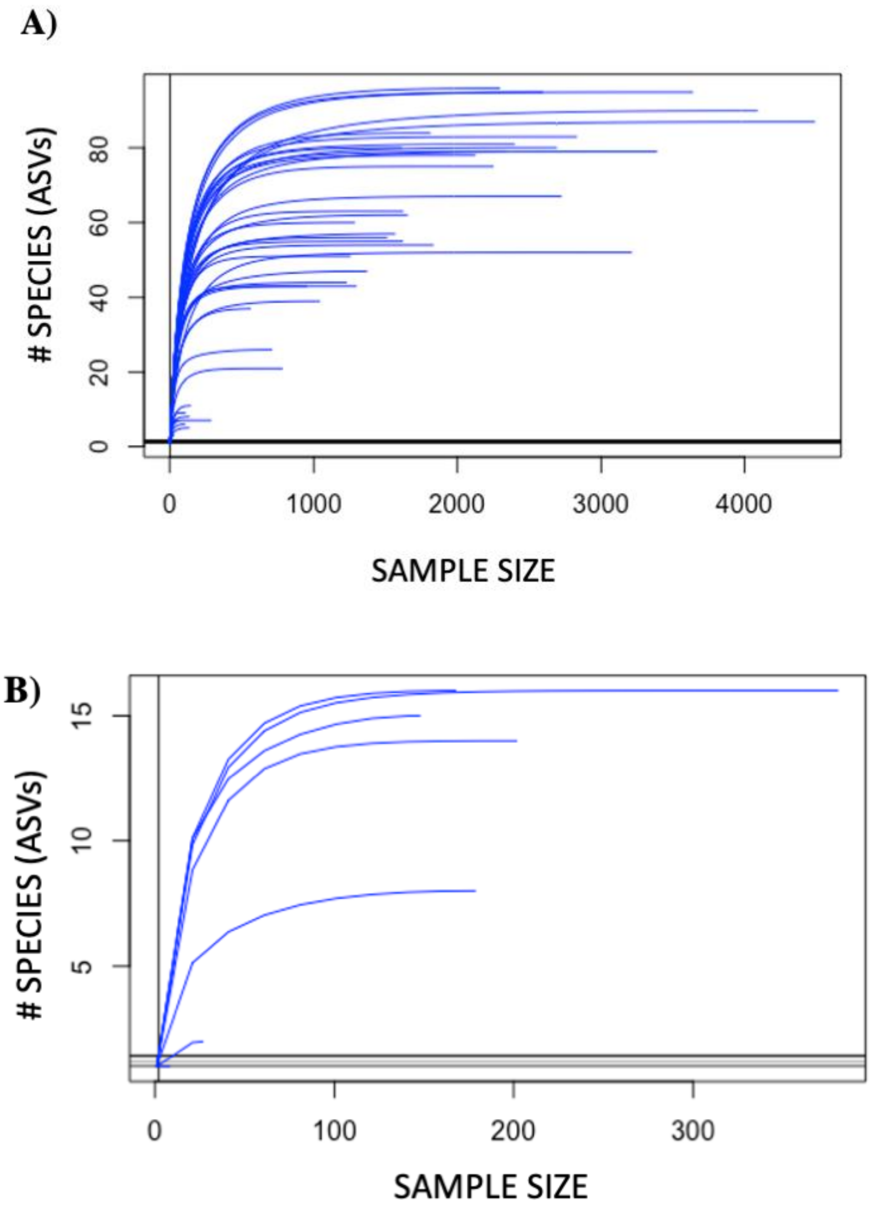


*Supplementary Figure 1*–Rarefaction curves of AE samples with <100 unique 18S ASVs (species) from the SPOT surface (A) and DCM (B). Five AE samples from the surface and two AE samples from the DCM were excluded from analyses due to insufficient diversity of 18S sequences. All samples with >100 unique 18S ASVs sufficiently captured the diversity of 18S organisms. Rarefaction curves were generated with the R package vegan version 2.5-6.

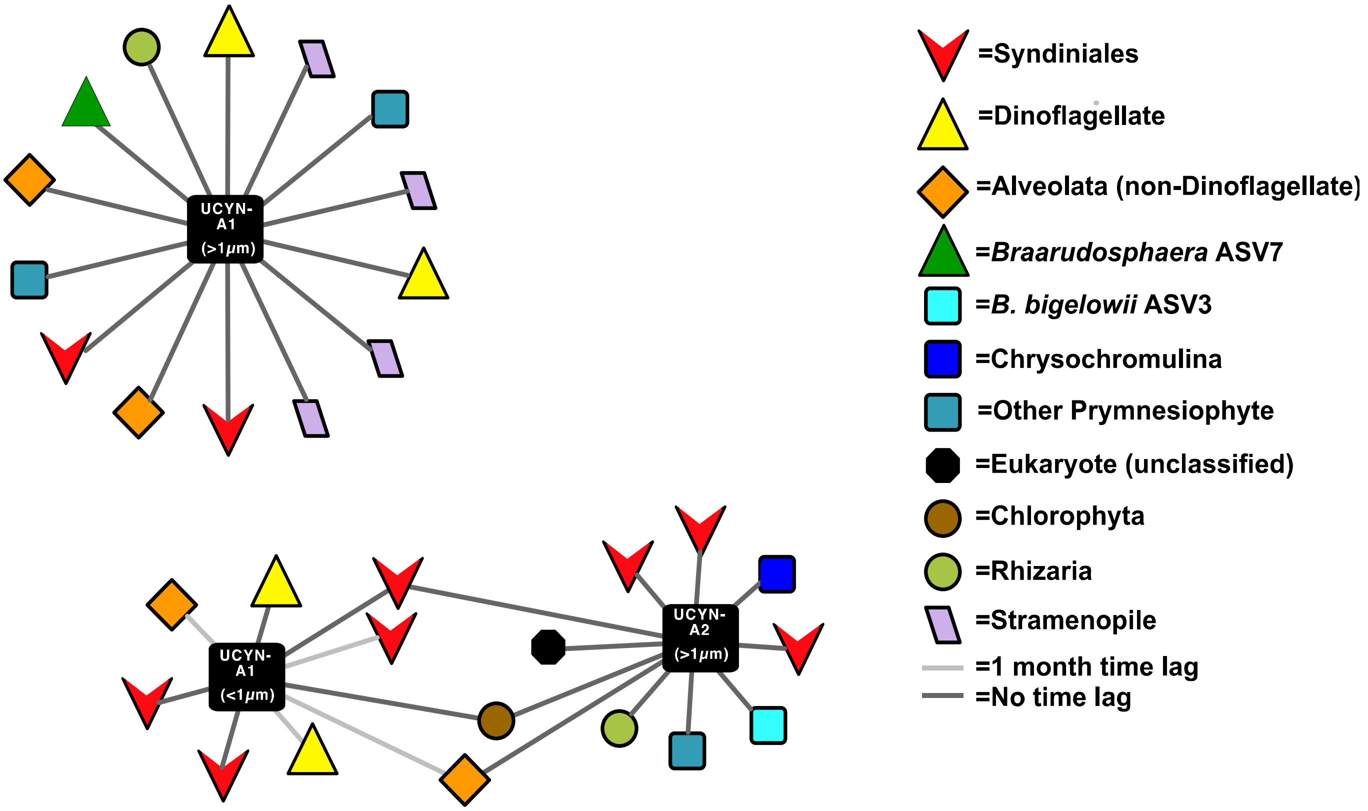


*Supplementary Figure 2*–eLSA networks constructed using interpolated, non-CLR transformed data from the SPOT surface (5m depth) miss interactions between UCYN-A and other taxa (compare to Figure 5). Networks were generated via eLSA and visualized in Cytoscape 3.5.

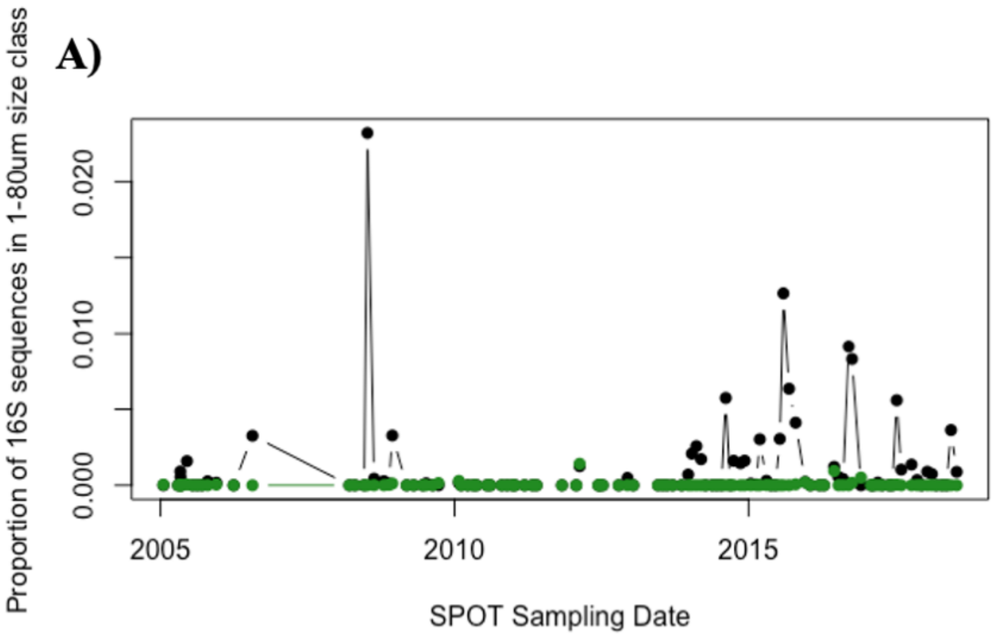
A)
Proportion of 16S sequences in 1-80um size class
0.000
0.010
0.020
2005
2010
2015
SPOT Sampling Date

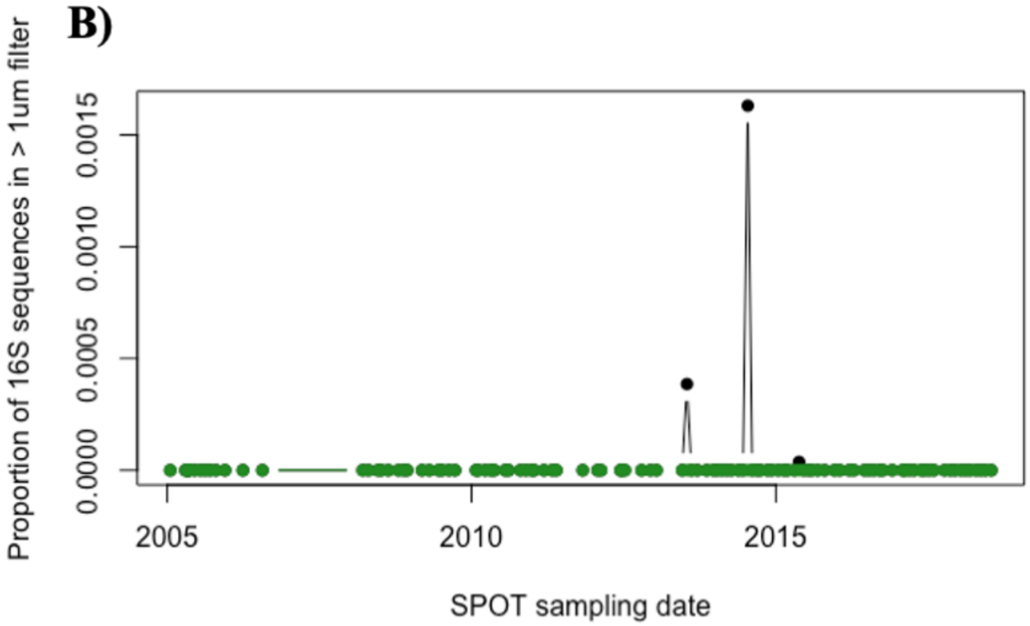
B)
Proportion of 16S sequences in > 1um filter
0.0000
0.0005
0.0010
0.0015
2005
2010
2015
SPOT sampling date

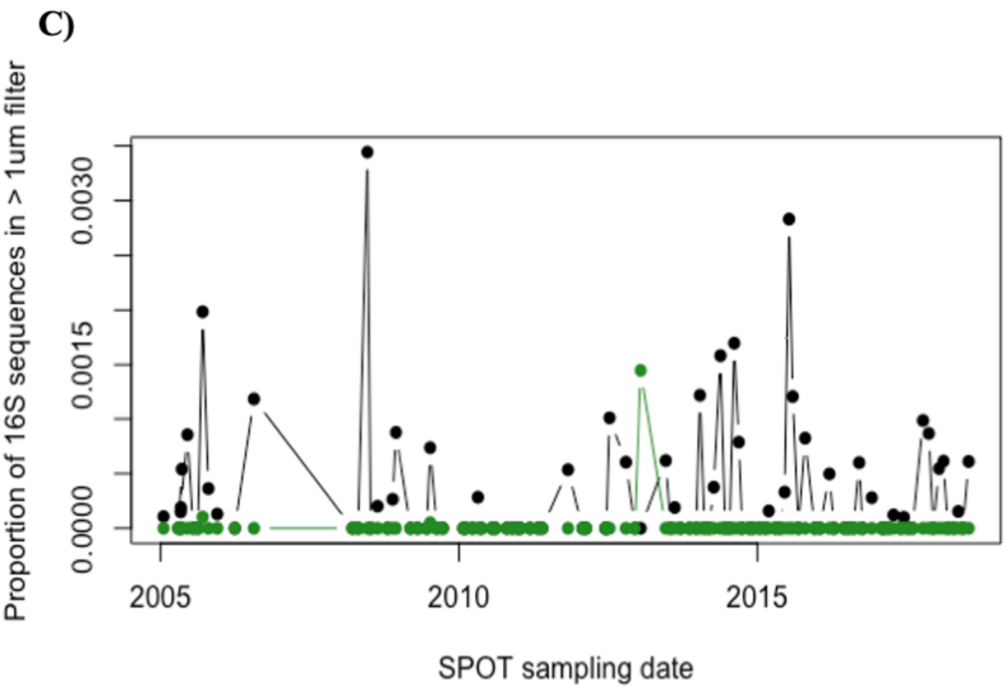


*Supplementary Figure 3*–Relative abundance of UCYN-A ASV1 (UCYN-A1 published genome) (A), UCYN-A ASV6 (B), and UCYN-A ASV 5 (UCYN-A2 published genome) (C) at the SPOT surface (black) and DCM (green) over time. Relative abundances are given out of the 16S community in the larger size fraction (AE filters; 1-80um).

*Supplementary Table 1*–Differences in environmental factors on dates UCYN-A1, UCYN-A2, hosts, and potential predator *Lepidodinium* ASVs are present vs. absent. Ekman transport is measured on the Cartesian coordinate system, such that movement Northwards and Eastwards is recorded as a more positive value. Positive MEI indicates El Niño events, while negative MEI represents La Nina conditions. P values were corrected for multiple testing via Benjamini-Hochberg correction. Boldface text indicates $p<0.05$, boldface, underlined text indicates $p<0.01$.

| | | UCYN-A1 (ASV1) | | | ***Braarudosphaera* ASV7x** | | | UCYN-A2 (ASV5) | | | ***B. bigelowii* ASV3** | | | | ***Lepidodinium* ASV** | | |
|---|---|---|---|---|---|---|---|---|---|---|---|---|---|---|---|---|---|
| | | Mean | Standard Error | P-values | Mean | Standard Error | P-values | Mean | Standard Error | P-values | Mean | Standard Error | P-values | | Mean | Standard Error | P-values |
| [NO2+NO3] (uM) | Present | 0.471 | 0.288 | 0.809 | 0.514 | 0.327 | 0.957 | 0.626 | 0.333 | 0.988 | 0.603 | 0.217 | 0.746 | Present | 0.620 | 0.355 | 0.754 |
| | Absent | 0.562 | 0.164 | | 0.531 | 0.152 | | 0.462 | 0.135 | | 0.494 | 0.260 | | Absent | 0.477 | 0.144 | |
| [PO4] (uM) | Present | 0.253 | 0.042 | 0.282 | 0.247 | 0.045 | 0.417 | 0.222 | 0.023 | 0.988 | 0.179 | 0.041 | 0.197 | Present | 0.246 | 0.048 | 0.754 |
| | Absent | 0.200 | 0.014 | | 0.208 | 0.017 | | 0.221 | 0.028 | | 0.239 | 0.011 | | Absent | 0.209 | 0.016 | |
| Bacterial Production (cells/mL/day) | Present | 284052.200 | 27744.452 | **0.006** | 305718.400 | 30237.819 | **0.050** | 365066.200 | 44642.812 | 0.988 | 329817.400 | 63965.889 | 0.197 | Present | 298758.800 | 23202.408 | 0.106 |
| | Absent | 481256.900 | 46408.102 | | 453865.400 | 44129.048 | | 422600.000 | 41673.916 | | 428844.800 | 22388.304 | | Absent | 451901.200 | 43959.033 | |
| MODIS [Chl] (mg * m-3) | Present | 0.507 | 0.070 | 0.350 | 0.451 | 0.046 | 0.124 | 0.509 | 0.051 | 0.988 | 0.444 | 0.377 | 0.197 | Present | 0.566 | 0.080 | 0.754 |
| | Absent | 0.901 | 0.286 | | 0.902 | 0.266 | | 0.883 | 0.277 | | 0.858 | 0.030 | | Absent | 0.827 | 0.255 | |
| MODIS SST (ºC) | Present | 19.090 | 0.400 | **0.004** | 19.079 | 0.431 | **0.025** | 18.878 | 0.347 | 0.446 | 19.242 | 0.415 | 0.059 | Present | 19.197 | 0.451 | **0.029** |
| | Absent | 17.604 | 0.247 | | 17.724 | 0.247 | | 17.796 | 0.292 | | 17.799 | 0.260 | | Absent | 17.714 | 0.241 | |
| CUTI index | Present | 0.326 | 0.019 | **0.004** | 0.345 | 0.021 | **0.025** | 0.416 | 0.026 | 0.988 | 0.360 | 0.033 | 0.197 | Present | 0.397 | 0.026 | 0.802 |
| | Absent | 0.458 | 0.023 | | 0.437 | 0.023 | | 0.396 | 0.022 | | 0.421 | 0.016 | | Absent | 0.407 | 0.022 | |
| BEUTI index | Present | 0.378 | 0.098 | **0.004** | 0.368 | 0.102 | **0.025** | 0.680 | 0.159 | 0.988 | 0.321 | 0.186 | 0.059 | Present | 0.630 | 0.163 | 0.719 |
| | Absent | 1.021 | 0.130 | | 0.977 | 0.124 | | 0.806 | 0.110 | | 0.933 | 0.057 | | Absent | 0.821 | 0.110 | |
| MEI | Present | -0.024 | 0.133 | **0.025** | -0.027 | 0.152 | **0.049** | -0.204 | 0.145 | 0.988 | -0.034 | 0.162 | 0.197 | Present | -0.183 | 0.160 | 0.754 |
| | Absent | -0.467 | 0.118 | | -0.431 | 0.110 | | -0.336 | 0.116 | | -0.386 | 0.114 | | Absent | -0.337 | 0.110 | |
| Magnitude of Surface Wind (m/s) | Present | 6.850 | 0.255 | **0.004** | 6.881 | 0.253 | **0.025** | 7.468 | 0.314 | 0.988 | 7.057 | 0.354 | 0.197 | Present | 7.284 | 0.297 | 0.718 |
| | Absent | 8.094 | 0.244 | | 7.980 | 0.243 | | 7.657 | 0.231 | | 7.796 | 0.199 | | Absent | 7.736 | 0.235 | |
| East-West Component of Ekman Transport (kg/m/s) | Present | -675.851 | 61.569 | **0.004** | -712.083 | 58.921 | **0.025** | -897.779 | 83.521 | 0.988 | -774.193 | 100.728 | 0.197 | Present | -815.777 | 70.719 | 0.719 |
| | Absent | -1039.232 | 71.392 | | -990.841 | 71.243 | | -885.622 | 65.968 | | -937.017 | 52.125 | | Absent | -928.030 | 68.823 | |
| North-South Component of Ekman Transport (kg/m/s) | Present | -512.139 | 84.007 | **0.004** | -536.981 | 80.845 | **0.025** | -767.567 | 101.137 | 0.988 | -583.699 | 119.591 | 0.197 | Present | -708.573 | 92.885 | 0.754 |
| | Absent | -919.728 | 82.514 | | -874.361 | 83.054 | | -743.356 | 79.260 | | -820.665 | 65.127 | | Absent | -775.010 | 81.202 | |

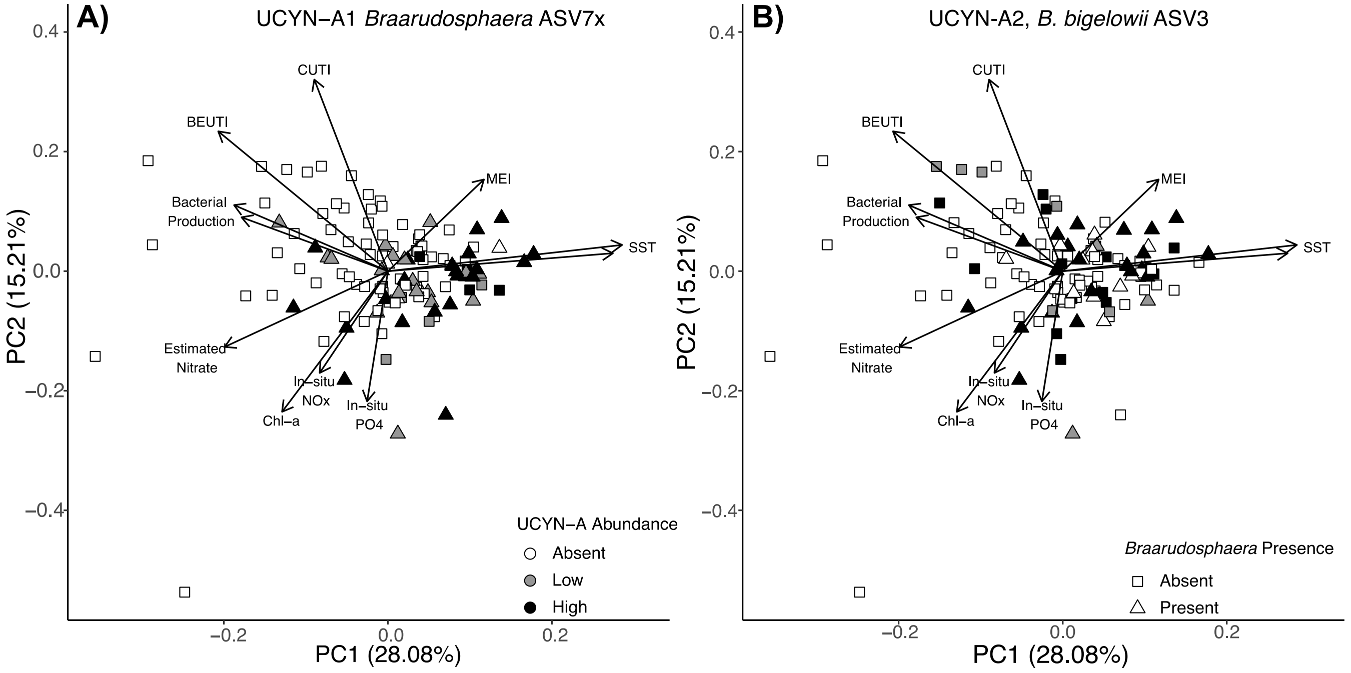


*Supplementary Figure 4*—Principal component analysis of environmental variables at the SPOT surface, overlaid with UCYN-A1 relative abundance and host presence/ absence (A) and UCYN-A2 relative abundance and host presence/ absence (B), show that temperature and MEI associate strongly with high relative abundance of UCYN-A1, but less strongly with that of UCYN-A2. UCYN-A ASVs were considered "absent" if they were <0.01% of the 16S community, "low abundance" if they were present in abundances lower than average (0.099% for ASV1, UCYN-A1, and 0.026% for ASV5, UCYN-A2), and "high abundance" if their relative abundance was greater than average. This is indicated by white, grey, and black points, respectively; host absence/ presence is indicated by squares vs. circles on both panels.

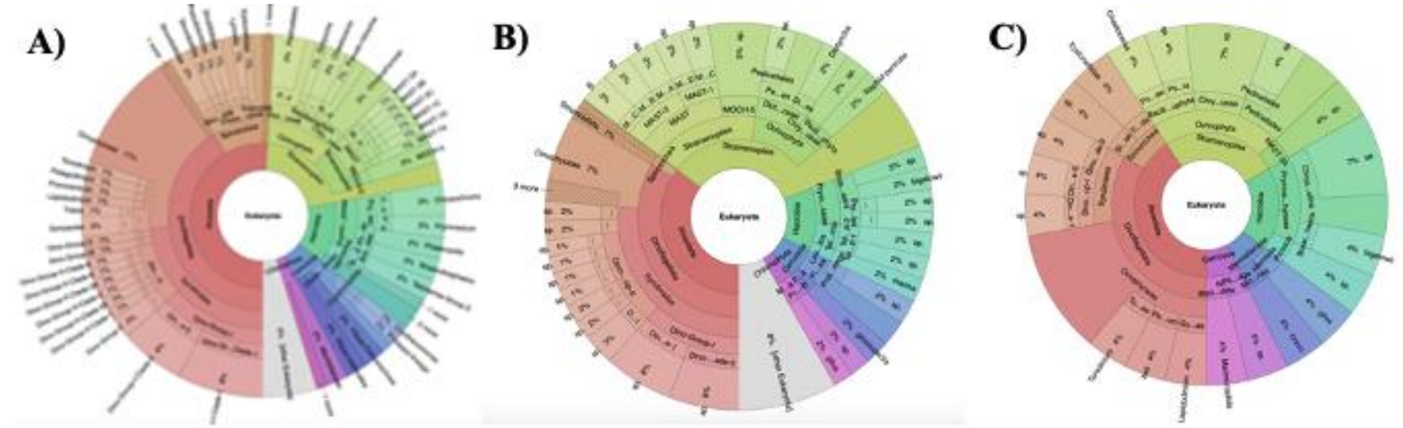


*Supplementary Figure 5*–Krona plot of 18S taxa that co-occur with A) all UCYN-A ASVs at SPOT surface, B) UCYN-A1 at the SPOT surface, C) UCYN-A2 at the SPOT surface. Co-occurrence data from the surface were filtered to a high level of statistical significance (P<0.005, Q<0.01).

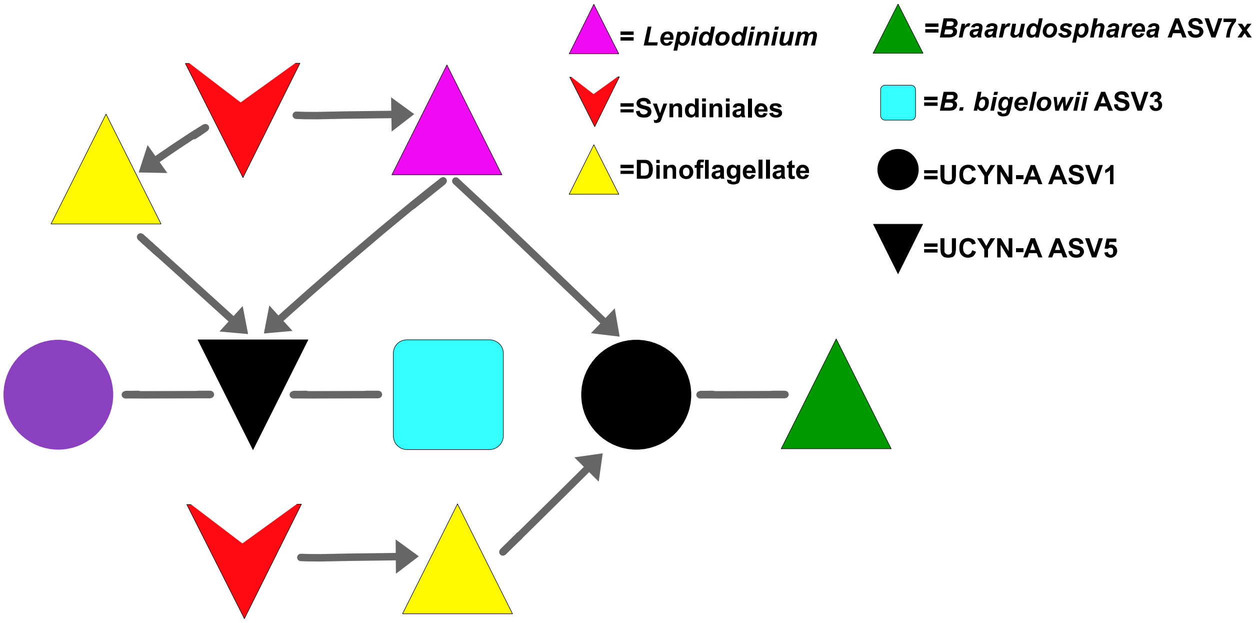


*Supplementary Figure 6*–Schematic of a hypothetical food web between UCYN-A ASVs and the 18S taxa with which they co-occur at the SPOT surface (see Figure 5). Arrows indicate predation/ parasitism; lines indicate symbiosis.

*Supplementary Table 2*–UCYN-A ASVs co-occur with a variety of 18S taxa. Hashes were generated via QIIME2; representative sequences were classified using SILVA132. Full 16S and 18S sequences are publicly available (see Data Availability Statement).

| Taxonomy | ASV Hash | Co-occurs with |
|---|---|---|
| Alveolata | 636f2ad7bbc9624ed6d8bd438d96f7b7 | UCYN-A1_0.22-1um |
| Alveolata | b930d2802540ff419424186646952b7d | UCYN-A1_0.22-1um |
| Alveolata | 3d3aaf7802f4db6f8bc887bb9e7774a8 | UCYN-A1_1-80um |
| Alveolata | 4b7935f69f43802ce9ea4b65a741845d | UCYN-A1_1-80um |
| Alveolata | 8e0bc96f64c6b0ca6e4aead874edee2a | UCYN-A1_1-80um |
| Alveolata | b6d8ad4a0a89749e0427b20d028f317d | UCYN-A1_1-80um |
| Alveolata | cccd609465491f29a68b22cdb1506556 | UCYN-A1_1-80um |
| Alveolata | 8a5ef38b1ab34ac0febbd7832a3ed0d1 | UCYN-A1_1-80um, UCYN-A1_0.22-1um |
| Alveolata | 2a77e539d5c3d4878fb0cc41c84a0fa3 | UCYN-A1_1-80um, UCYN-A1_0.22-1um, UCYN-A2 |
| Alveolata | a9666cf5e782f71ad780c73f0f59c119 | UCYN-A2 |
| Braarudosphaera_sp_7x | be3cdecefbceb0d8b25a2e42ed058b50 | UCYN-A1_1-80um, UCYN-A1_0.22-1um |
| Braarudospharea_bigelowii_3 | 70a5283da28db501a349c5beb22881e7 | UCYN-A1_1-80um, UCYN-A2 |
| Chlorophyta | fd7bca02dc31e29cb36624e39cbcc27a | UCYN-A1_1-80um, UCYN-A1_0.22-1um |
| Chlorophyta | c3b3a5a8e14ea2c9cdb58028bc55bca0 | UCYN-A1_1-80um, UCYN-A1_0.22-1um, UCYN-A2 |
| Chrysochromulina | d55992e6da65321a9b3c0ce3426e73ac | UCYN-A1_1-80um, UCYN-A2 |
| Chrysochromulina | eaaf40a3c970e0ec2167de48c4b001eb | UCYN-A2 |
| Dinoflagellate | 26cc4edeb2a1788fc8fdf147923c06b4 | UCYN-A1_0.22-1um |
| Dinoflagellate | 997d8cc0ee56b8f185884eb47f7bbbd6 | UCYN-A1_0.22-1um |
| Dinoflagellate | 4bb7bd73a0c0f6a55e0c4acecdbfd099 | UCYN-A1_1-80um |
| Dinoflagellate | 70241d94140d1eaea54e19bc52b1bdd1 | UCYN-A1_1-80um |
| Dinoflagellate | 791b0b5d59992db5031a18510b325ced | UCYN-A1_1-80um |
| Dinoflagellate | ead0d51affbef7dd17d17e483eb0f244 | UCYN-A1_1-80um |
| Dinoflagellate | ef4dcdee685b3fd3427dff66b3a9e379 | UCYN-A1_1-80um |
| Dinoflagellate | f383ee229e9cae5c76c76062e8a0f99f | UCYN-A1_1-80um |
| Dinoflagellate | a2bcca80cacedb46e9e55cd424022684 | UCYN-A1_1-80um, UCYN-A2 |
| Dinoflagellate | dc228f26735fb8483e2e7a8f07994c23 | UCYN-A1_1-80um, UCYN-A2 |
| Dinoflagellate | 23f2bbf2ecbae94d2b57c207aa8abc62 | UCYN-A2 |
| Dinoflagellate | 95e7881e06d651005303f80a0824f7f9 | UCYN-A2 |
| Dinoflagellate | eaee3f386d9d2d2c4ea3d90ccf81f48a | UCYN-A2 |
| Dinoflagellate | ec296e5b3180d5c65f9e81e7165a4763 | UCYN-A2 |
| Eukaryote | 0edea9a57e97085c27800904bc55437d | UCYN-A1_1-80um |
| Eukaryote | af7721ace95e845243e2b715dbcca683 | UCYN-A1_1-80um |
| Eukaryote | 72ad5bae980d6f3a205494139435e4e5 | UCYN-A1_1-80um, UCYN-A1_0.22-1um |
| Lepidodinium | c114523e0bef5840b096693e46f441a2 | UCYN-A1_1-80um, UCYN-A1_0.22-1um, UCYN-A2 |
| Prymnesiophyte | 23206df8663c546b75a92fadb5de9b33 | UCYN-A1_0.22-1um |
| Prymnesiophyte | 79c87d3c4954a1dec1972fc679befcb5 | UCYN-A1_1-80um |
| Prymnesiophyte | 98d1369a93242f295deda7ee996b9886 | UCYN-A1_1-80um |
| Prymnesiophyte | 803aa95deeb6167672a24a67029f2b82 | UCYN-A1_1-80um, UCYN-A1_0.22-1um |
| Prymnesiophyte | ffa92571884bcbe0193ce4a1e4e843be | UCYN-A1_1-80um, UCYN-A1_0.22-1um |
| Prymnesiophyte | 55bc25d102de6079cc48ba48515e72e2 | UCYN-A2 |
| Rhizaria | 307e50f0be54fdc6ac0ea8183ff77d0e | UCYN-A1_0.22-1um |
| Rhizaria | a615f49d7d2fbed5b85069fb94087fa3 | UCYN-A1_1-80um |
| Rhizaria | ded52caa99b40106f449ecc43dbbeee0 | UCYN-A1_1-80um |
| Rhizaria | 4f2711ffbb0611ea359dd96680df1c4f | UCYN-A2 |
| Rhizaria | afacfa595bc8d424f2a648990538e0d4 | UCYN-A2 |
| Stramenopile | eac29d65650c1a316e01b9fa8a5a9038 | UCYN-A1_0.22-1um |
| Stramenopile | 25e0c3351c5b93e80e0f02e6ba23c077 | UCYN-A1_1-80um |
| Stramenopile | 3465aaadf57e9cf04c930da3beb3deef | UCYN-A1_1-80um |
| Stramenopile | 5ca13089ea80484bd62871929d00bf95 | UCYN-A1_1-80um |
| Stramenopile | 6f39e2bbe0e4c6fa1e2eb082348de7c5 | UCYN-A1_1-80um |
| Stramenopile | 8940d219ac059e1421b8910843b4fd82 | UCYN-A1_1-80um |
| Stramenopile | 9ec148397bce05c40a8d97a91574a1ef | UCYN-A1_1-80um |
| Stramenopile | ddaade5c44a6bbd3a74f879c85b5faf3 | UCYN-A1_1-80um |
| Stramenopile | e67e2f1a5016412cf9826b35523352c4 | UCYN-A1_1-80um |
| Stramenopile | e966740cb469db7493a9b384262bce67 | UCYN-A1_1-80um |
| Stramenopile | a94291b249e004b97f399d41c5cc4b82 | UCYN-A1_1-80um, UCYN-A1_0.22-1um |
| Stramenopile | 5d86bdd765a6ce520791f28dda5819ed | UCYN-A1_1-80um, UCYN-A2 |
| Stramenopile | 3d03bbd7eb3a06f19d2f377b5eb2efb5 | UCYN-A2 |
| Stramenopile | e09f07091bd33255025bd0a1cec414ca | UCYN-A2 |
| Syndiniales | 00399bcfe64d51dac93b62a40832b514 | UCYN-A1_0.22-1um |
| Syndiniales | 10657154c17634d9006a00243a3736b1 | UCYN-A1_0.22-1um |
| Syndiniales | 38caa165f588a30638f74c2edb93662c | UCYN-A1_0.22-1um |
| Syndiniales | 69e6b69c7b0addcc5c9c44d08015e45b | UCYN-A1_0.22-1um |
| Syndiniales | bce9eb5a6281547d9fffcd95d9156913 | UCYN-A1_0.22-1um |
| Syndiniales | 1082ec476ee33ebf0800a137e9dd6730 | UCYN-A1_1-80um |
| Syndiniales | 12516469ae16dbe47afd1383f1a38d29 | UCYN-A1_1-80um |
| Syndiniales | 43d72e28390647a43a479322275ebabc | UCYN-A1_1-80um |
| Syndiniales | 4f0a7afd789e6f14b745c17acad17068 | UCYN-A1_1-80um |
| Syndiniales | 6fa9b449505f9b00f18edbedca877d17 | UCYN-A1_1-80um |
| Syndiniales | 80d903e0cb7271e69b2ec8bd1f87f45c | UCYN-A1_1-80um |
| Syndiniales | b22089acf5e02fe3477be0ae8fe10cb5 | UCYN-A1_1-80um |
| Syndiniales | bf238b09c39ad60be96eb93f30ac6915 | UCYN-A1_1-80um |
| Syndiniales | eefc3152825b60051b3f78c504aca2a9 | UCYN-A1_1-80um |
| Syndiniales | f14396a0e7efc97700da47bcb853bfad | UCYN-A1_1-80um |
| Syndiniales | 365026beab40dc3dacd12678ac56cfc2 | UCYN-A1_1-80um, UCYN-A1_0.22-1um |
| Syndiniales | 6bbbadb60359057c81abc84e7d0b797a | UCYN-A1_1-80um, UCYN-A1_0.22-1um |
| Syndiniales | b3a109d089b15d7d926497af27f7fd2c | UCYN-A1_1-80um, UCYN-A1_0.22-1um |
| Syndiniales | 624af6018837349e853bc2e772461e80 | UCYN-A1_1-80um, UCYN-A1_0.22-1um, UCYN-A2 |
| Syndiniales | 68216c70c4cc7ae340a7f17cb6973f5b | UCYN-A1_1-80um, UCYN-A1_0.22-1um, UCYN-A2 |
| Syndiniales | 6a30306b1212b48152ee097506fedad2 | UCYN-A2 |
| Syndiniales | 6c3205e48cb5c1cc405609e73063607d | UCYN-A2 |
| Syndiniales | 6cf6cfba7e7b46309c2fc38b44a28064 | UCYN-A2 |

*Supplementary Figure 7*–Relative abundances of UCYN-A 16S sequences linearly correlate with relative abundances of host 18S sequences. A) Pairwise comparison of all UCYN-A vs. all *Braarudospharea* ASVs in the SPOT dataset. B) UCYN-A1 (ASV1) and UCYN-A2 (ASV5) relative abundances correlate with one another, hence these symbionts appear to correlate with one another's host organisms. Relative abundances are given out of the whole microbial community (16S + 18S) in the 1-80um size fraction at 5m depth.

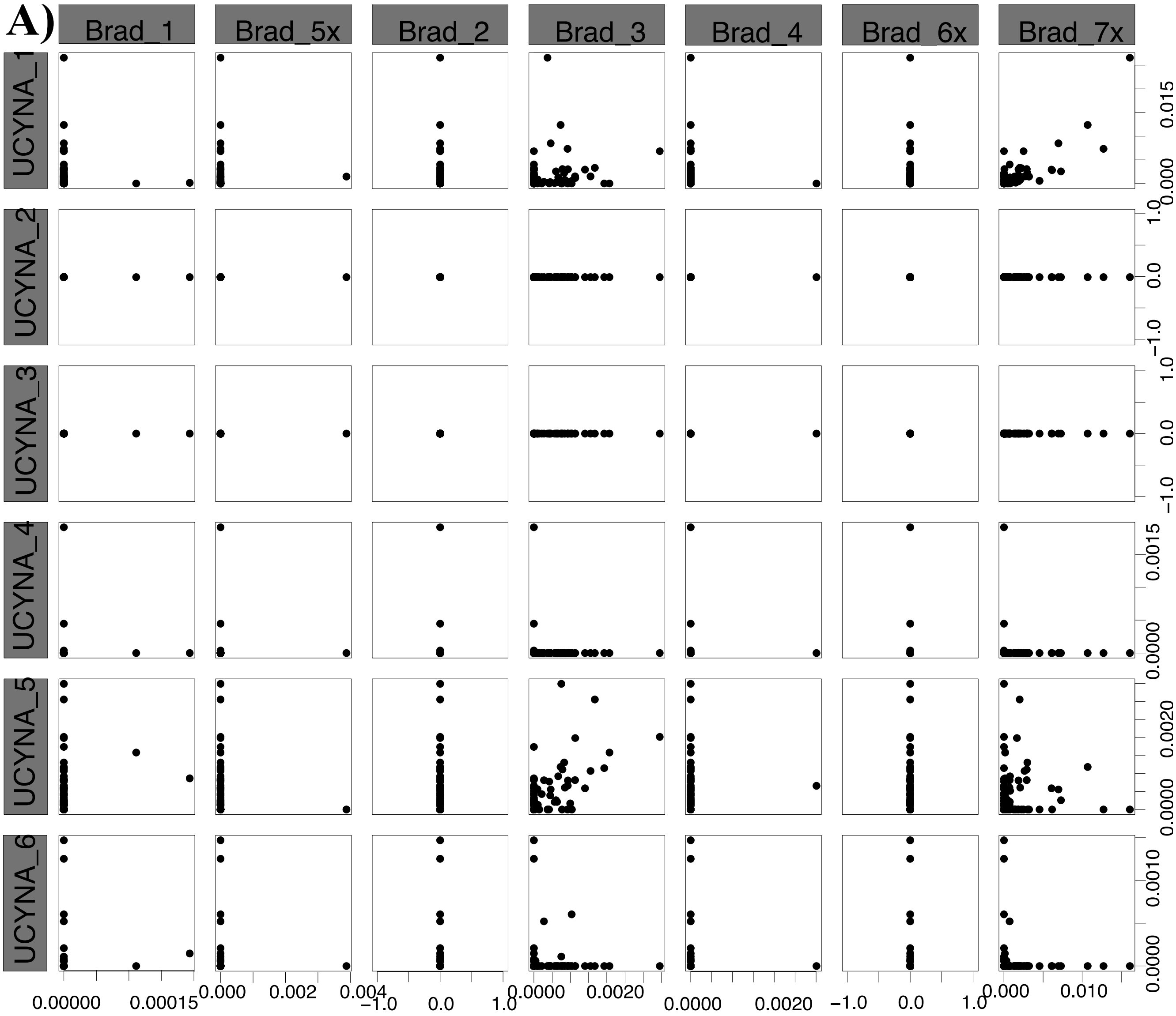

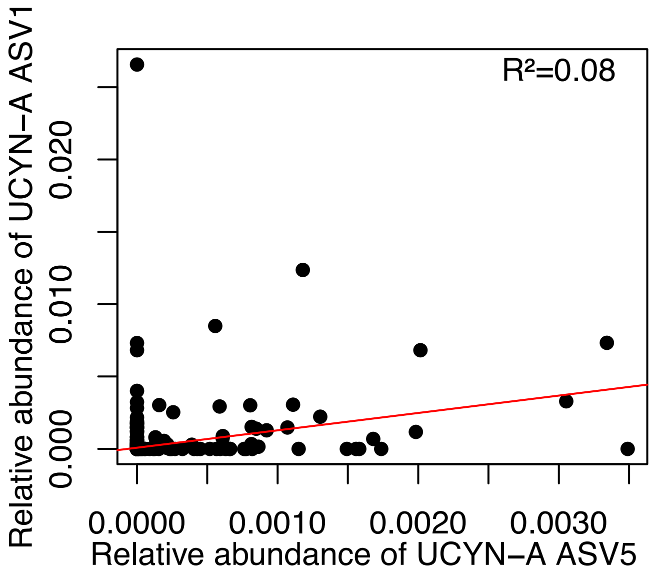
R²=0.08
Relative abundance of UCYN−A ASV1
0.000
0.010
0.020
0.0000
0.0010
0.0020
0.0030
Relative abundance of UCYN−A ASV5

B)

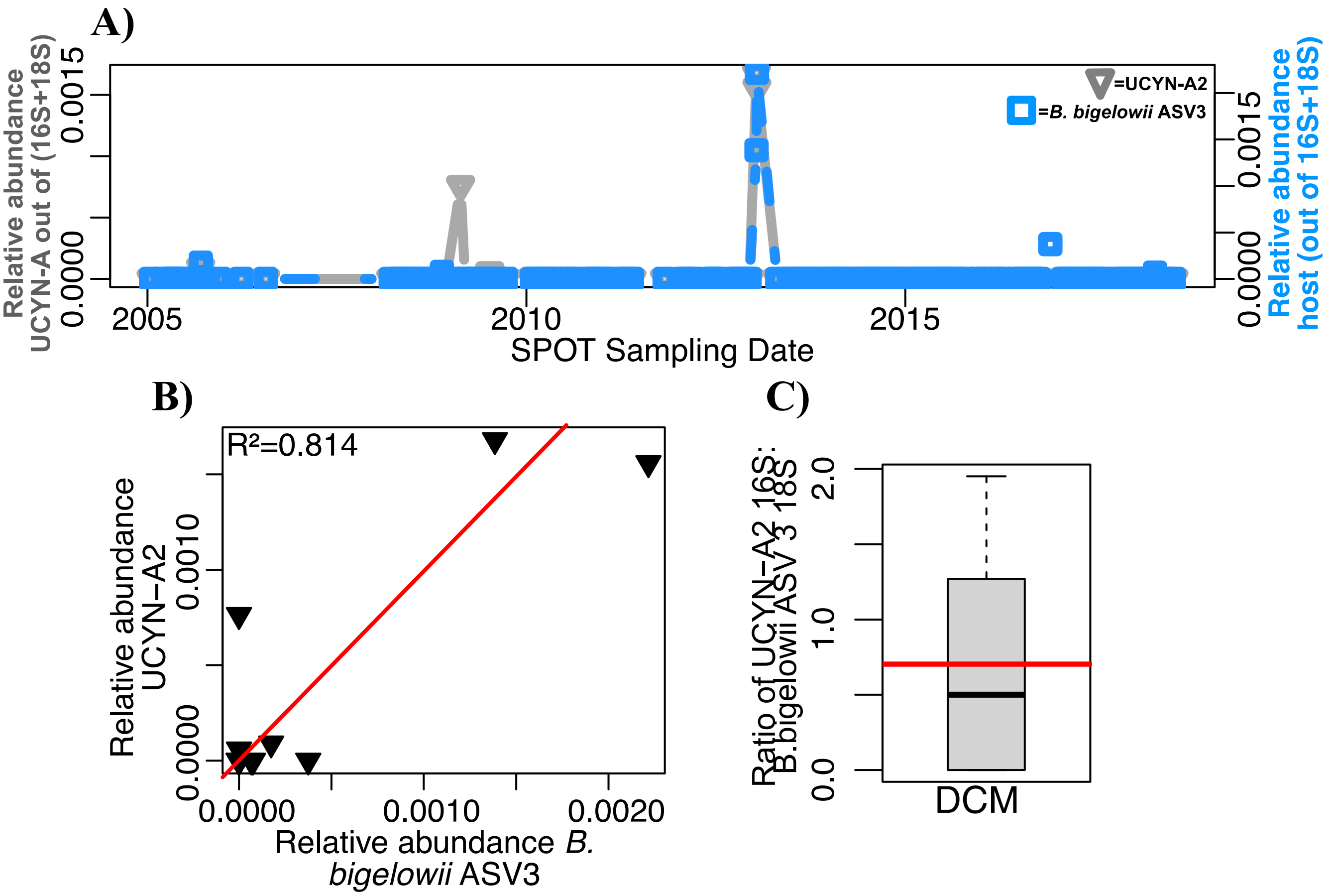
A)
Relative abundance UCYN-A out of (16S+18S)
0.0000
0.0015
=UCYN-A2
=B. bigelowii ASV3
Relative abundance host (out of 16S+18S)
2005
2010
2015
SPOT Sampling Date
B)
R²=0.814
Relative abundance UCYN−A2
0.0000
0.0010
0.0020
Relative abundance B. bigelowii ASV3
C)
Ratio of UCYN−A2 16S: B.bigelowii ASV 3 18S
0.0
1.0
2.0
DCM

*Supplementary Figure 8*–A) UCYN-A2 co-occurs with its most common host at the DCM across the SPOT time series. B) UCYN-A2 relative abundance correlates with its putative host relative abundance at the DCM. C) The ratio of 16S: 18S genes of these organisms is, on average, as expected. Boxplot values indicate the median and IQR values of this ratio; the red line indicates the average (0.703).

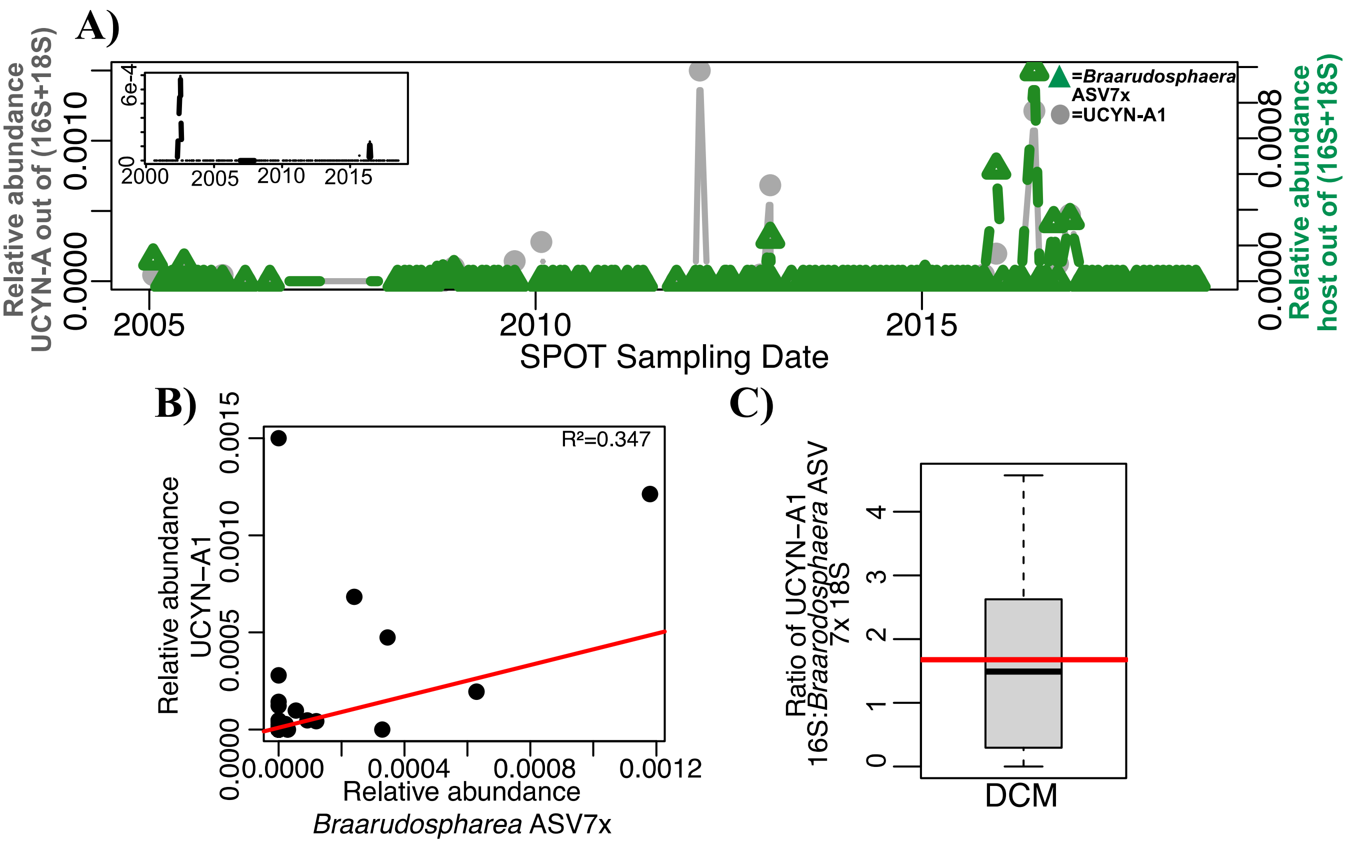


*Supplementary Figure 9*–A) UCYN-A1 co-occurs with its putative host at the DCM across the SPOT time series. Panel inset indicates the relative abundance of UCYN-A1 in the smaller size fraction. B) UCYN-A1 relative abundance correlates with its putative host relative abundance. C) The ratio of 16S: 18S genes of these organisms is, on average, as expected. Boxplot values indicate the median and IQR values of this ratio; the red line indicates the average (1.674).

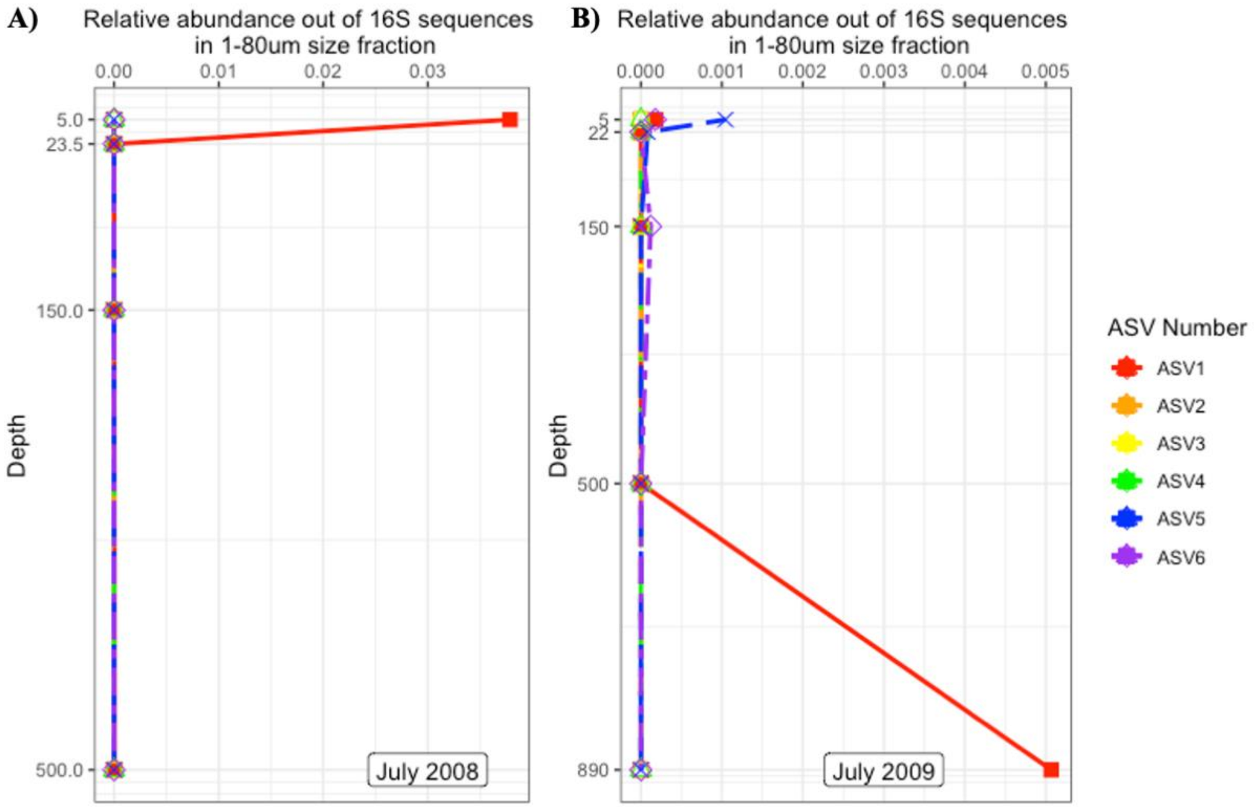


*Supplementary Figure 10*–Relative abundance of UCYN-A ASVs in the larger size fraction over depth in A) July 2008, the date UCYN-A1 reached its maximum relative abundance, and B) July 2009, the date UCYN-A1 appeared at 890m.

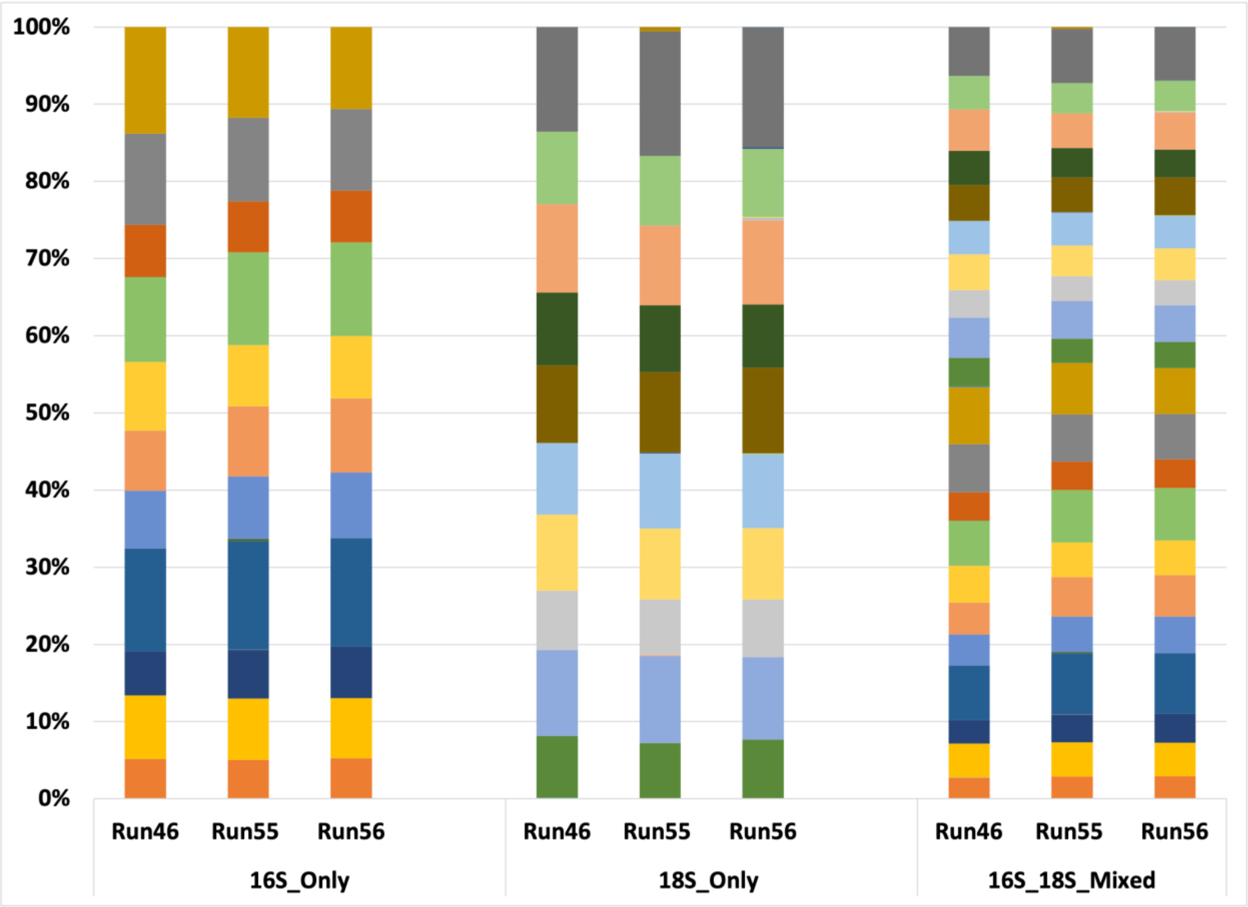


*Supplementary Figure 11*–Mixed mock communities contain even proportions of 16S (right) and 18S (middle) sequences. DNA from the small subunit of the rRNA gene of 21 organisms was pooled and sequenced in equal concentrations on an Illumina HiSeq or MiSeq. Normalized mixed mock communities match the proportions of sequenced DNA, indicating normalization was successful (left).

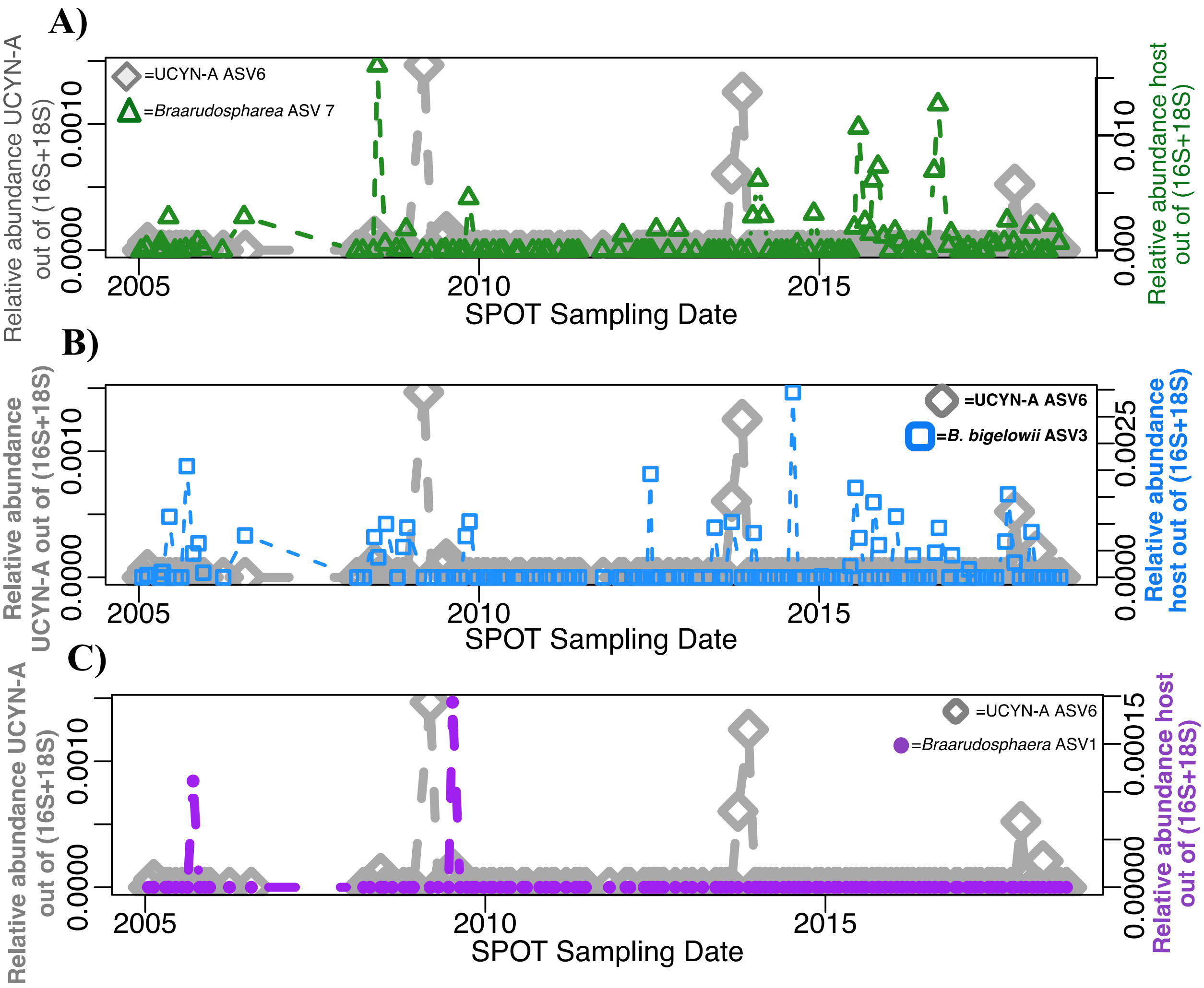


*Supplementary Figure 12*–UCYN-A ASV6 does not co-occur with any *Braarudosphaera* ASV across the SPOT dataset at 5m.
