## Supplementary material for "Symbiotic diazotrophic UCYN-A strains co-occurred with El Niño, relaxed upwelling, and varied eukaryotes over 10 years off Southern California Bight": Interactive HTML file with 18S taxa that co-occur with all UCYN-A ASVs: SupplementaryFigure5A_18S_wall_UCYNA_ASVs_krona.html

Javascript must be enabled to view this page.

magnitude
score


Euk\_taxa

 104

 5.04807692307692

 104

 5.04807692307692

 53

-.905660377358491

 1

 1

 1

 1

 1

 1

 1

 1

 1

 1

 9

 2

 9

 2

 3

 2

 1

 2

 1

 2

 2

 2

 1

 2

 4

 2

 1

 2

 1

 2

 2

 2

 2

 2

 1

 2

 1

 2

 2

 2

 1

 2

 43

-1.55813953488372

 18

-1

 3

-1

 1

-1

 1

-1

 2

-1

 2

-1

 1

-1

 1

-1

 1

-1

 1

-1

 1

-1

 1

-1

 1

-1

 1

-1

 1

-1

 1

-1

 1

-1

 1

-1

 1

-1

 1

-1

 1

-1

 1

-1

 24

-2

 14

-2

 8

-2

 8

-2

 6

-2

 6

-2

 8

-2

 2

-2

 2

-2

 1

-2

 1

-2

 1

-2

 1

-2

 1

-2

 1

-2

 1

-2

 1

-2

 1

-2

 1

-2

 1

-2

 1

-2

 1

-2

 1

-2

 1

-2

 1

-2

 1

-2

 1

-2

 3

 3

 3

 3

 2

 3

 2

 3

 2

 3

 1

 3

 1

 3

 1

 3

 1

 3

 1

 3

 1

 3

 14

 12.2142857142857

 9

 16

 9

 16

 2

 16

 2

 16

 2

 16

 5

 16

 3

 16

 3

 16

 2

 16

 2

 16

 2

 16

 2

 16

 2

 16

 1

 4

 1

 4

 1

 4

 1

 4

 1

 4

 1

 5

 1

 5

 1

 5

 1

 5

 1

 5

 3

 6

 3

 6

 3

 6

 1

 6

 1

 6

 2

 6

 2

 6

 7

 8.14285714285714

 7

 8.14285714285714

 2

 7

 1

 7

 1

 7

 1

 7

 1

 7

 1

 7

 1

 7

 2

 8

 1

 8

 1

 8

 1

 8

 1

 8

 1

 8

 3

 9

 2

 9

 1

 9

 1

 9

 1

 9

 1

 9

 22

 12.0909090909091

 12

 11.6666666666667

 4

 10

 4

 10

 2

 10

 2

 10

 2

 10

 1

 10

 3

 15

 3

 15

 3

 15

 3

 15

 5

 11

 5

 11

 1

 11

 1

 11

 4

 11

 3

 11

 8

 12.25

 6

 12

 2

 12

 1

 12

 1

 12

 1

 12

 1

 12

 3

 12

 1

 12

 1

 12

 1

 12

 1

 12

 1

 12

 1

 12

 1

 12

 1

 12

 1

 12

 2

 13

 2

 13

 2

 13

 2

 13
