## Supplementary material for "Symbiotic diazotrophic UCYN-A strains co-occurred with El Niño, relaxed upwelling, and varied eukaryotes over 10 years off Southern California Bight": Interactive HTML file with 18S taxa that co-occur with UCYN-A1: SupplementaryFigure5B_18S_w_UCYNA_ASV1_krona.html

Javascript must be enabled to view this page.

magnitude
score


A1\_data

 2556

 5.44561815336463

 2556

 5.44561815336463

 903

-1.59579180509413

 1

 1

 1

 1

 1

 1

 1

 1

 1

 1

 1

 1

 44

 2

 44

 2

 9

 2

 2

 2

 2

 2

 2

 2

 7

 2

 3

 2

 3

 2

 26

 2

 5

 2

 5

 2

 5

 2

 13

 2

 13

 2

 13

 2

 8

 2

 8

 2

 8

 2

 9

 2

 858

-1.78321678321678

 186

-1

 21

-1

 21

-1

 21

-1

 10

-1

 11

-1

 12

-1

 12

-1

 12

-1

 13

-1

 13

-1

 13

-1

 13

-1

 672

-2

 297

-2

 147

-2

 147

-2

 147

-2

 150

-2

 150

-2

 150

-2

 292

-2

 67

-2

 67

-2

 67

-2

 35

-2

 35

-2

 35

-2

 36

-2

 36

-2

 36

-2

 37

-2

 37

-2

 37

-2

 38

-2

 38

-2

 38

-2

 39

-2

 39

-2

 39

-2

 40

-2

 40

-2

 40

-2

 41

-2

 41

-2

 41

-2

 41

-2

 42

-2

 42

-2

 42

-2

 42

-2

 87

 3

 87

 3

 43

 3

 43

 3

 43

 3

 43

 3

 43

 3

 44

 3

 44

 3

 44

 3

 44

 3

 44

 3

 336

 16

 186

 16

 186

 16

 91

 16

 45

 16

 45

 16

 45

 16

 46

 16

 46

 16

 46

 16

 95

 16

 95

 16

 95

 16

 47

 16

 48

 16

 49

 16

 49

 16

 49

 16

 49

 16

 49

 16

 49

 16

 101

 16

 101

 16

 101

 16

 50

 16

 50

 16

 50

 16

 51

 16

 51

 16

 51

 16

 159

 7

 159

 7

 52

 7

 52

 7

 52

 7

 52

 7

 52

 7

 107

 7

 53

 7

 53

 7

 53

 7

 53

 7

 861

 10

 285

 10

 55

 10

 55

 10

 55

 10

 56

 10

 56

 10

 56

 10

 56

 10

 56

 10

 174

 10

 174

 10

 57

 10

 57

 10

 117

 10

 58

 10

 58

 10

 441

 10

 310

 10

 121

 10

 60

 10

 60

 10

 60

 10

 61

 10

 61

 10

 61

 10

 189

 10

 62

 10

 62

 10

 62

 10

 63

 10

 63

 10

 63

 10

 64

 10

 64

 10

 64

 10

 131

 10

 131

 10

 131

 10

 131

 10

 131

 10
