## Supplementary material for "Symbiotic diazotrophic UCYN-A strains co-occurred with El Niño, relaxed upwelling, and varied eukaryotes over 10 years off Southern California Bight": Interactive HTML file with 18S taxa that co-occur with UCYN-A2: SupplementaryFigure5C_18S_wUCYNA2_krona.html

Javascript must be enabled to view this page.

magnitude
score


A5\_data

 27

 6.11111111111111

 27

 6.11111111111111

 11

-.727272727272727

 2

 2

 2

 2

 1

 2

 1

 2

 1

 2

 1

 2

 1

 2

 1

 2

 9

-1.33333333333333

 6

-1

 1

-1

 1

-1

 1

-1

 1

-1

 1

-1

 1

-1

 1

-1

 1

-1

 1

-1

 1

-1

 3

-2

 2

-2

 1

-2

 1

-2

 1

-2

 1

-2

 1

-2

 1

-2

 1

-2

 1

-2

 1

-2

 1

-2

 2

 3

 2

 3

 2

 3

 2

 3

 2

 3

 1

 3

 1

 3

 1

 3

 1

 3

 5

 13.8

 4

 16

 4

 16

 3

 16

 3

 16

 3

 16

 2

 16

 1

 16

 1

 16

 1

 16

 1

 16

 1

 5

 1

 5

 1

 5

 1

 5

 1

 5

 1

 5

 2

 7

 2

 7

 1

 7

 1

 7

 1

 7

 1

 7

 1

 7

 1

 7

 1

 7

 1

 7

 7

 12

 6

 12

 2

 10

 2

 10

 1

 10

 1

 10

 1

 10

 1

 10

 1

 10

 2

 15

 2

 15

 2

 15

 2

 15

 2

 15

 2

 11

 2

 11

 2

 11

 1

 11

 1

 11

 1

 12

 1

 12

 1

 12

 1

 12

 1

 12

 1

 12
